# Humanized Klotho haplotypes cause widespread transcriptomic changes in mouse brain

**DOI:** 10.64898/2026.04.15.718745

**Authors:** Anna L. Tyler, Dylan Garceau, Kevin P. Kotredes, Annat Haber, Catrina Spruce, Ravi S. Pandey, Christoph Preuss, Michael Sasner, Gregory W. Carter

## Abstract

Klotho *KL* is an aging factor that has been associated with Alzheimer’s Disease (AD) risk. Two common alleles circulate in human populations: the major allele FC and the minor allele VS, which is defined by two SNPs that cause two amino acid substitutions (F352V and C370S) in *KL*’s second exon. To investigate the possibility that human *KL* variants influence brain aging and cognition, we developed a novel mouse model with humanized *KL* alleles. We used RNA-Seq to measure the whole brain transcriptome in four-and 12-month-old male and female C57Bl/6J mice carrying either the FC or the VS *KL* allele. We found that FC and VS carriers had widespread differences in gene expression in the brain at 12 months old, but not at four months old. The largest differences were in genes annotated to mitochondrial, ribosomal, and synaptic functions. Differential exon usage analysis identified differential splicing of synaptic genes, further supporting a role for *KL* on neuronal function. A more focused analysis of differential expression identified variation in glutamate receptors and amyloid precursor (APP) processing in particular, thereby linking human *KL* haplotypes to biological processes integral to AD pathogenesis. These results provide evidence that the human FC and VS *KL* haplotypes affect the function of the *KL* protein product in a manner that has widespread effects on gene expression in the brain and supports the hypothesis that these haplotypes may influence AD risk and pathogenesis.

## Introduction

Klotho (*KL*, also called 𝛼-*KL*) is a gene encoding a transmembrane protein that is expressed in the kidney, choroid plexus in the brain, and liver and has been associated with longevity ^1^. Loss-of-function mutations in *Kl* in mice causes marked progeria including many hallmarks of aging such as osteoporosis, atherosclerosis, skin atrophy, and death by two months of age ^2^. This phenotype is 100% penetrant and consistent across multiple strain backgrounds ^2^. Conversely, overexpression of *Kl* extends lifespan, particularly in male mice ^3^. In addition to its generic effects on aging, *Kl* appears to have effects on cognition. Genetic overexpression of *Kl* in mice enhances cognitive performance ^4^, and injection of KL protein to Rhesus macaques enhances learning and memory for weeks after administration ^5^.

Due to its profound effects on the aging process and on cognition, *Kl* is considered a promising target for modulating age-related cognitive decline in diseases such as Alzheimer’s Disease (AD). Promising results have been produced in experiments in which overexpression of *Kl* in mice with humanized amyloid precursor protein (*APP*) improves both AD-related pathology and cognition ^6^.

Multiple genetic studies support the idea that genetic variation in *KL* is related to lifespan and cognition in humans as well. Initial studies identified two major haplotypes of *KL* defined by a two-SNP haplotype. These SNPs, rs9536314 and rs9527025, cause two amino acid substitutions in exon 2 of the human gene (F352V and C370S). The two resulting haplotypes are named for these amino acid substitutions and are called FC and VS. The KL-FC haplotype is the more common with an allele frequency of 87% ^7^, although allele frequency varies widely across populations from almost 0% in East Asians to 20% in Ashkenazi Jews ^7^. Global minor allele (VS) frequency is around 13% ^7^.

Initial epidemiological studies associated the VS allele with increased longevity in a Czech population ^8^. Consistent with this observation, *in vitro* studies suggested that the VS allele resulted in a 1.6-fold increase in secretion of the *Kl* protein ^9^. A second study in an independent population supported the longevity advantage of individuals heterozygous for the VS allele (with apparent disadvantage for VS homozygotes) ^10^. The increased longevity was accompanied by reduced risk factors for cardiovascular disease and stroke ^10^.

Research since these original associations has had mixed results. Multiple studies have found association between the VS allele and improved aging outcomes, such as increased longevity ^8;10–13^, reduced cardiovascu-lar disease ^10;12^, improved cognition ^10;14^, and reduced risk of AD ^15;16^ and Parkinson’s disease ^17^. However, others have found either no association between the VS allele and healthy aging ^18^ and cognition ^12;19;20^, or a reversed relationship in which the VS allele was associated with increased risk for coronary artery disease ^21^ or cognitive decline ^22^.

One potential source of the mixed results in humans are the many sources of variation across human populations, such as variation in genetic background and environmental conditions. KL levels vary significantly with diet ^23;24^, exercise ^25^, stress levels ^26^ and time of day ^27^, so human studies may not be able to detect genotype-dependent differences in expression. Cohort studies are confounded by systematic variation across age groups that can only be addressed by longitudinal studies, which are difiicult in humans.

To better study the precise effects of the FC and VS alleles, we used inbred mice as a model of human aging and cognition. The effects of KL depletion and overexpression have been well characterized in mice ^2;3;6^. The genetic models described here introduced the human *KL* variants to mice allowing us to study their effects on the brain transcriptome, under controlled conditions. Here we introduced the human VS and FC *KL* alleles into male and female C57BL/6J mice. We compared these to wild type B6 mice which carry the FS haplotype, a combination of the two SNPs not observed in human populations. We used bulk RNA-Seq to measure transcription in brain at four months and 12 months of age to investigate age-related effects of these alleles on brain transcription.

## Methods

### Mouse model development

All experiments were approved by the Animal Care and Use Committee at The Jackson Laboratory. Mice were bred in the mouse facility at The Jackson Laboratory and maintained in a 12/12-h light/dark cycle, consisting of 12 h-ON 7 am-7 pm, followed by 12 h-OFF. Room temperatures are maintained at 18-24^∘^C (65-75^∘^F) with 40-60% humidity. All mice were housed in positive, individually ventilated cages (PIV). Standard autoclaved 6% fat diet (Purina Lab Diet 5K52) was available to the mice *ad libitum*, as was water with acidity regulated from pH 2.5-3.0. Novel mouse alleles were generated using direct delivery of CRISPR-Cas9 reagents to B6J (C57BL/6J; JAX #664) mouse zygotes. Analysis of genomic DNA sequence surrounding the target region, using the Benchling (www.benchling.com) guide RNA design tool, identified appropriate gRNA sequences with a suitable target endonuclease site.

**KL-FC mice** carry the S370C point variant (rs9527025) in exon 2 of the Klotho gene, homologous to the human late-onset Alzheimer’s disease “risk” configuration of p.F352 and p.C370. The CRISPR guide 5’-GACTTTTTTGCTCTCTCCTT-3’ was used to cut at the desired mutation site, and the repair oligonu-cleotide 5’-CCAAGACAGAAGCTGCCTCAGGTTGGGAGACTCCAATTGGCGGAACTTCATGTTAGG GTCCAATAGCTGAAAGCTCAAGGTTGGTCCGAAG**<u>C</u>**A**<u>A</u>**AGAGCAAAAAAGTCAGCAGTTCCTC TGATGAGC-3’ was used to introduce the S370C variant (TCC>TGC) as well as an upstream silent mutation immediately 5’ (CTC>CTT) to prevent re-cutting.

**KL-VS mice** carry the F352V variant (rs9536314) in exon 2 of the Klotho gene, homologous to the human late-onset Alzheimer’s disease “neutral” configuration of p.V352 and p.S370. The CRISPR guide 5’-ATTTTACTGAATCTGAGAAG-3’ was used to cut at the desired mutation site, and the repair oligonu-cleotide 5- GGAACTTCATGTTAGGGTCCAATAGCTGAAAGCTCAAGGTTGGTCCGAAGGAGAGAG CAAAAAAGTCAGCAGTTCCTCTGATGAG**<u>G</u>**CTCTTCTCAGATTCAGTAA**<u>C</u>**ATCAGGCAGAAGAG ACGAGAG-3’ was used to introduce the F352V variant (TTT>GTT) as well as a silent mutation (AGG>AGC) just downstream to prevent re-cutting.

For each model, PCR products spanning the entire donor region were sequenced to verify the appropriate sequence (see Figure 1 and Supplemental Figure 1). Founders were backcrossed to B6J for at least three generations prior to intercrossing to create homozygotes.

**Figure 1:**
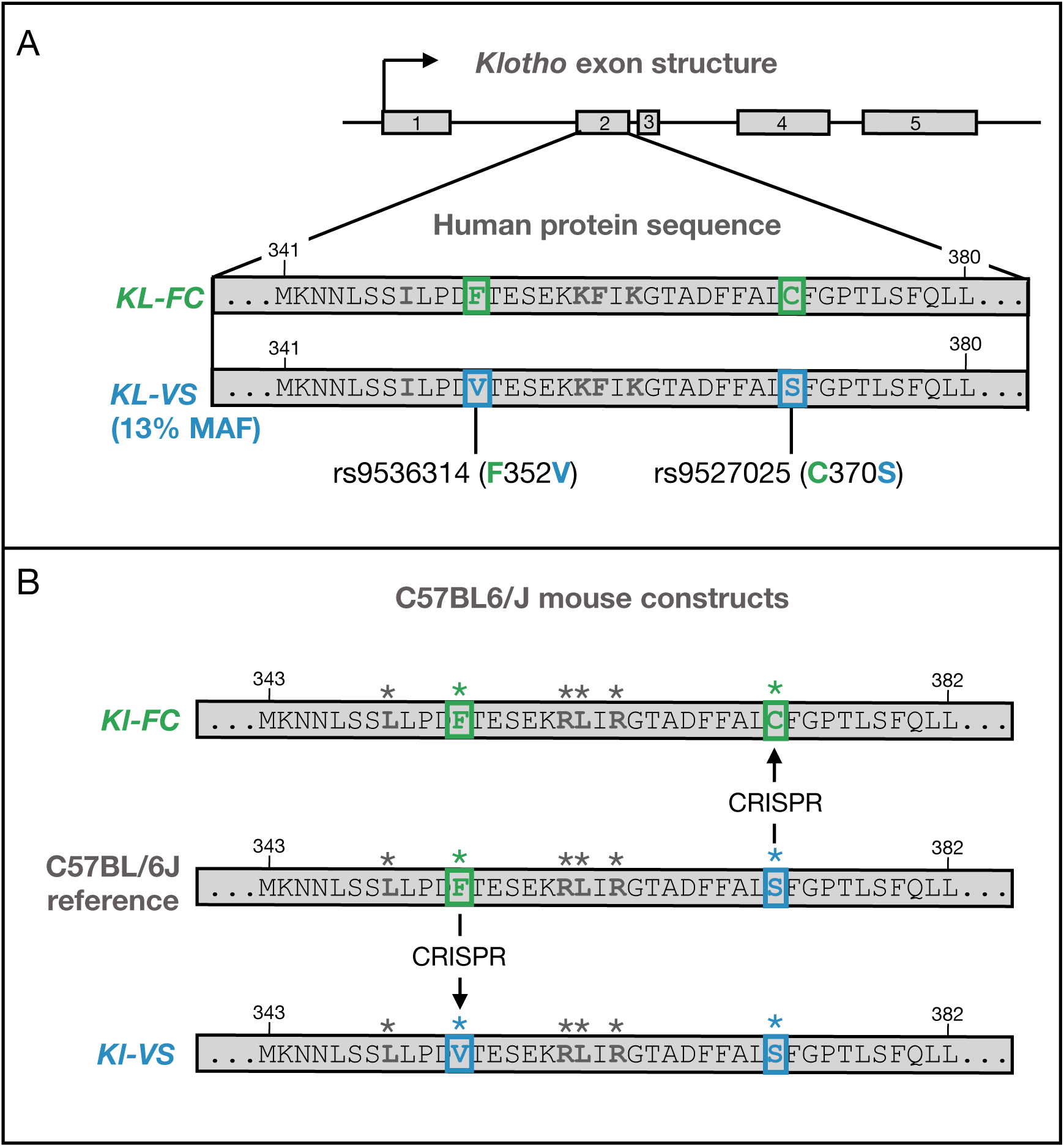
*KL*/*Kl* structure and mouse constructs. **A.** The *KL* gene has five exons. The second exon (expanded) contains two common SNPs (rs9536314 and rs9527025), which cause two amino acid changes (F352V and C370S respectively). These two SNPs are represented in two common human haplotypes, the *KL*-FC haplotype is the major allele (frequency 87and the *KL*-VS haplotype is the minor allele (frequency 13**B.** The C57BL6/J mouse version of *Kl* differs from the common human haplotype at the C372S (human C370S) position rendering it different from either human haplotype. We used CRISPR to introduce each human SNP into the mouse allele creating a *KL*-FC allele and a *KL*-VS allele to compare to the wild type mouse allele *Kl*-FS. Asterisks indicate residues that differ in the human protein.

The KL-FC model (S370C) is available from the Jackson Lab as C57BL/6J-Kl^em1Adiuj^/J (JAX #36534). The KL-VS model (F352V) is available as C57BL/6J-Kl^em2Adiuj^/J (JAX #36535). Allele-specific genotyping protocols are available on JAX mice data sheets for each model.

The number of mice in each group is detailed in Table 1.

**Table 1:** Number of mice in each group.

| haplotype | Female |  | Male |  |
| --- | --- | --- | --- | --- |
|  | 4 mo. | 12 mo. | 4 mo. | 12 mo. |
| WT/WT | 5 | 10 | 8 | 9 |
| WT/FC | 5 | 4 | 5 | 6 |
| FC/FC | 3 | 4 | 5 | 6 |
| WT/VS | 2 | 5 | 2 | 3 |
| VS/VS | 2 | 5 | 4 | 6 |

### Sample Collection

Brain samples were collected from mice four- and 12-months old. For each of age cohort, four to six mice for each sex, genotype and timepoint were chosen. Hemi-brains were dissected and snap frozen for gene expression. RNA-sequencing was performed at The Jackson Laboratory (Bar Harbor, ME).

### Animal anesthesia

Upon arrival at the terminal endpoint for each aged mouse cohort, individual animals were weighed prior to intraperitoneal administration of either: (A) ketamine (100mg/kg) and xylazine (10mg/kg); or (B) tribromoethanol (1mg/kg). Routine confirmation of deep anesthesia was performed every five minutes by toe pinch.

### Whole animal perfusion

First confirming deep anesthetization via toe pinch, animals are secured to a surgical board or tray using needles or pins and abdomen wetted with 70% ethanol followed by an incision along the ventral midline along the entire ventral surface, exposing the underlying muscle of the thorax and abdomen. An additional incision is made into this underlying muscle and cut to puncture the diaphragm, taking care not to cut any major blood vessels or the lungs. To expose the heart the ribcage can be cut along the lateral borders and removed. A small incision is made in the right atrium of the heart to relieve diastolic pressure and begin perfusing the animal transcardially with 1xPBS. To clear the vascular system of all blood a butterfly catheter needle is inserted into the left ventricle attached to a perfusion pump. Approximately 10mL of 1xPBS solution will clear the system of a 20g animal. Completion of perfusion and clearance of the vascular system was indicated by a blanching of the liver and PBS will be noticed exiting the right atrium.

### Brain collection

Anesthetized and subsequently perfused animals were decapitated, and heads submerged quickly in cold 1xPBS. The skin was cut from the base of the neck, over the top of the skull, between the ears, stopping between the eyes, and separated to either side to expose the skull. The skull was cut dorsally until the cerebellum. Two opposite, horizontal cuts were made in the skull under, but without cutting, the cerebellum to ease separation of the skull at the midline. On the top of the skull, scissor blades were inserted superficially at the bregma and slowly expanded to separate the skull down the midline. With a pair of blunt-end forceps, the two skull plates were removed to expose the brain. The brain was carefully removed from the skull, weighed, and divided midsagitally, into left and right hemispheres, using a brain matrix. The right hemisphere was quickly homogenized by chopping and mixing with razor blades (∼50 strokes), on stainless steel stage, on ice, and equally aliquoted into three cryotubes transcriptomic analysis. Cryotubes were immediately snap frozen on liquid nitrogen, and stored long-term at -80^∘^C.

### RNA Sample Extraction

Total RNA from fresh frozen tissues were isolated using QIAGEN RNeasy Mini Kit according to the manufacturer’s instructions. Briefly, the tissue (up to 30 mg) was homogenized in Buffer RLT, supplemented with 1% B-mercaptoethanol using Bead Bug with a speed set at 2500 g and 20 seconds per time (up to three times for homogenization). Equal volume of 70% ethanol was added to the lysate and transferred to the RNeasy spin column. After centrifugation, the flow-through was discarded. This was followed by on-column

DNase I treatment. 80 uL of DNase I mix (70 uL Buffer RDD with 10 uL DNase I) was added directly to the RNA spin column, incubated for 10 minutes at room temperature, and then washed. Finally, total RNA was eluted in 50 𝜇L RNase-free water, and stored at -80^∘^sC. RNA concentration and quality were assessed using the Qubit RNA BR Assay (Thermo Scientific) and the RNA ScreenTape (Agilent Technologies), respectively, according to the manufacturers’ instructions.

### RNA Library Preparation and Sequencing

RNA-seq libraries were prepared with KAPA mRNA Hyperprep kit (Roche) according to manufacturer’s instruction. RNA samples were normalized to 300 ng input in total 44 𝜇L and 6 𝜇L of 1:1000 dilution ERCC (mix 1 or mix 2) (Thermo Fisher) was spiked in to each sample. First, poly A RNA was isolated from total RNA using oligo-dT magnetic beads. Purified RNA was then fragmented at 85^∘^C for six mins, targeting fragments range 250-300 bp. Fragmented RNA is reverse transcribed with an incubation of 25^∘^C for 10 mins, 42^∘^C for 15 mins and an inactivation step at 70^∘^C for 15mins. This was followed by second strand synthesis and A-tailing at 16^∘^C for 30 mins and 62^∘^C for 10 min. A-tailed, double stranded cDNA fragments were ligated with Illumina unique adaptors (Illumina). Adaptor-ligated DNA was purified using Ampure XP beads. This is followed by 10 cycles of PCR amplification. The final library was cleaned up using AMpure XP beads. Library concentration and quality were assessed using the Qubit HS dsDNA Assay (Thermo Scientific) and the D5000 ScreenTape (Agilent Technologies), respectively, according to the manufacturers’ instructions.

### Sequencing

The Illumina libraries were normalized and pooled. Quantification of library pool was performed using real-time qPCR (KAPA and Thermo Fisher). The final library pool was normalized to 2 nM. The pool was then denatured and loaded on the Illumina sequencer as per the manufacturer’s instructions (Illumina). PhiX was spiked in at 1%. Sequencing was performed on Illumina NovaSeq 6000 platform generating paired end reads of 151 bp.

### RNA Data Processing and Analysis

RNA-Seq data were processed using nf-core/rnaseq pipeline [https://doi.org/10.5281/zenodo.1400710]. Briefly, reads were aligned to the reference mouse genome (version GRCm38.p97) using STAR and gene expression was quantified with RSEM. Genotypes were validated using samtools mpileup to count nucleotide identity of reference vs alternate alleles at the FC and VS locations. Sex was validated by comparing expression of the sex-specific genes *Xist* (ENSMUSG00000086503), *Eif2s3y* (ENSMUSG00000069049), and *Ddx3y* (ENSMUSG00000069045).

### Differential expression

We used vst normalization in the R package DESeq2^28^ to normalize gene read counts and regressed out the effects of sequencing batch.

We used multiple linear regression to investigate the effects of sex, age, and *KL* haplotype on the residuals of each gene. We used the following linear model to look for main effects and interaction effects of each variable as a factor.

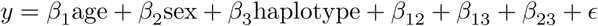

Because there was no interaction between sex and *KL* status on gene expression in our initial analysis, we regressed sex out of expression for all subsequent analyses.

## Differential exon usage

We used the R package DRIMSeq ^29^ to analyze differential exon usage in the bulk RNA-Seq data. To maximize power to detect variation, we compared the FC carriers to the VS carriers.

We filtered exons to those expressed in at least 10 individuals and having an average of at least 10 reads across samples. We used sequencing batch and sex as additive covariates and used an adjusted 𝑝 value ≤ 0.05 as a significance threshold.

### Biological domains and subdomains

Previous work curated 19 biological domains (Biodomains) and 80 subdomains that are associated with AD-related processes including synaptic function, APP metabolism, immune response, and lipid metabolism ^30;31^. Each domain was defined using sets of gene ontology (GO) terms selected to be coherent within each domain but independent of the other domains. The 19 Biodomains have since been further broken down into 80 biological subdomains for more specific classification of genomic patterns ^31^. These domains and subdomains were designed to identify AD-related biological processes involved in large-scale variation in transcriptomic patterns.

AD Biodomain gene lists and associated GO Terms were obtained from the Alzheimer’s Disease Knowledge Portal under syn25428992 (https://adknowledgeportal.synapse.org/FileEntity?entityId=syn25428992) on October 25, 2024.

We calculated a mean expression for the genes in each subdomain in all individual mice, thereby defining the average subdomain expression for each mouse.

We then fit a linear model to explain subdomain mean expression using *KL* haplotype as the explanatory factor:

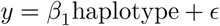

where 𝑦 is the mean expression for the subdomain, and haplotype is an ordered factor with the following levels: *KL*-FC < *KL*-WT < *KL*-VS. For both the FC and VS haplotypes, the heterozygous and homozygous animals had similar expression (Supplemental Figure 2). Because of this similarity, we grouped the heterozygotes and homozygotes of each haplotype.

We used the variance explained by this model as the strength of the relationship between *KL* haplotype and biological subdomain gene expression.

### KEGG pathways

The Kyoto Encyclopedia of Genes and Genomes (KEGG) ^32;33^ is a curated database of genomic data organized for bioinformatic analysis of cellular and higher level functions of biological organisms. One highly used category in this database is the collection of pathways, which are expertly curated maps that represent known molecular interactions and reactions in cellular pathways. These pathways include well-understood pathways such as glycolysis (map00010) and oxidative phosphorylation (map00190), as well as higher level functional relationships among genes, such as those involved in neurodegeneration (map05022) and cancer (map05200).

We used the R package clusterProfiler ^34^, to download KEGG pathways on October 20, 2024.

### Human-Mouse Orthologs

We used the R package BioMart ^35;36^ to match mouse-human orthologs.

### KEGG-Biodomain Intersections

We calculated the subset of genes in the intersections of all pairs of KEGG pathways and Biodomains. For each intersection containing at least 10 genes, we calculated the mean expression of all genes across all individuals and concatenated these means into a single matrix in which rows were KEGG-Biodomain intersections, and columns were individual mice.

For each KEGG-Biodomain intersection (each row of the matrix), we fit a linear model as described above to assess the effect of *KL* haplotype on mean gene expression of the intersectional genes. We selected intersections with an FDR ≤ 0.001 as significant. This threshold was selected to be sufiiciently stringent to control false positives while including a reasonable number of pathway intersections for visualization. Higher thresholds did not change any of the patterns in the results, and mostly introduced more redundancy. We used the effect of haplotype from each model as a useful way to sort intersections. Positive effects corresponded to upregulation in VS animals relative to FC and WT, and negative effects corresponded to downregulation in VS animals relative to FC and WT.

### Comparison to human iPSC-derived brain organoids

We compared the effects of *KL* haplotype in the mouse brain to the transcriptomic effects of *KL* expression induction in human induced pluripotent stem cell (iPSC) cortical brain organoids ^37^. Although different from human brain tissue, organoids offer an accessible, experimentally manipulable system in which to observe the effects of simple interventions on complex conglomerations of human neuronal cells. In this case, the authors measured the transcriptomic effects of a 1.5-fold induced increase of expression of endogenous *KL* in cortical brain organoids. We downloaded RNA-Seq read counts from the Gene Expression Omnibus ^38;39^ (https://www.ncbi.nlm.nih.gov/gds/) accession number GSE171719.

We filtered transcripts to those with at least 10 counts across all samples and used use the variance stabiliza-tion transformation (vst) in the R package DESeq2^28^. We grouped genes based on the KEGG-Biodomain intersections described above and calculated mean expression of each gene across the three replicates in each group (*KL*-induced and untreated). To compare the effects of *KL* overexpression directly to the effects of haplotype in the mice, we fit linear models to describe mean gene expression in each group using treatment as a factor. We compared the 𝛽 coefiicient for treatment to the 𝛽 coefiicient for the *KL* haplotype in the mice.

### Data Availability

**AD Biodomain gene lists** are available on the Alzheimer’s Disease Knowledge Portal under syn25428992 https://www.synapse.org/Synapse:syn25428992.

**Mouse expression data** and metadata will be made available on the Alzheimer’s Disease Knowledge Portal upon publication.

**Organoid expression** data are available at the Gene Expression Omnibus, accession number GSE171719 https://www.ncbi.nlm.nih.gov/geo/query/acc.cgi?acc=GSE171719.

### Code Availability

All code used for analyses and figure generation are available on GitHub: https://github.com/annaLtyler/ klotho_analysis.

## Results

### Human *KL* haplotypes were introduced into C57BL/6J mice

To investigate the potential of the *KL*-VS allele to alter AD pathology and clinical outcomes, we generated mouse models carrying the two humanized *KL* haplotypes (Figure 1A). The C57BL/6J (B6) mouse KL protein has 86% amino acid identity with the human protein. However, it differs from the human reference genome at position C372S (C370S in humans) such that the B6 haplotype (*KL*-FS) is a mixture of the two human haplotypes (*KL*-FC and *KL*-VS). We used CRISPR to introduce the human SNPs into the mouse *Kl* gene to create two humanized haplotypes, *KL*-FC and *KL*-VS. We compared these haplotypes to the wildtype mouse haplotype, *Kl*-FS. There were no overt phenotypic differences among the *KL* haplotypes. Transcriptomic differences are detailed below.

### *KL* haplotype was not associated with variation in *KL* mRNA abundance in the brain

In brain tissue, the FC and VS alleles of *KL* were not associated with differential mRNA levels of *KL* itself (linear regression: 𝑛_𝐹𝐶_ = 44, 𝑛_𝑊𝑇_ = 43, 𝑛_𝑉 𝑆_ = 45, 𝛽 = -0.025, 𝑅^2^ = -0.012, 𝑝 = 0.82) (Fig. 2A). Studies in human subjects have suggested that the VS allele may be associated with greater 𝛼-KL protein levels in serum ^40;41^ and CSF ^41^, however, there are multiple control points between mRNA production and protein levels that may result in differential protein levels independent of mRNA expression. Further, other organs, like the kidney are primary sources of serum KL, and it is possible that transcription of *KL* in these tissues, varied between the haplotypes. *KL* gene expression also did not significantly vary by sex (linear regression: 𝑛_𝐹_ = 60, 𝑛_𝑀_ = 72, 𝛽 = -0.0054, 𝑅^2^ = -0.0077, 𝑝 = 0.98) or age (linear regression: 𝑛_4𝑚_ = 64, 𝑛_12𝑚_ = 68, 𝛽 = 0.18, 𝑅^2^ = 0.0086, 𝑝 = 0.15) (Fig. 2A).

**Figure 2:**
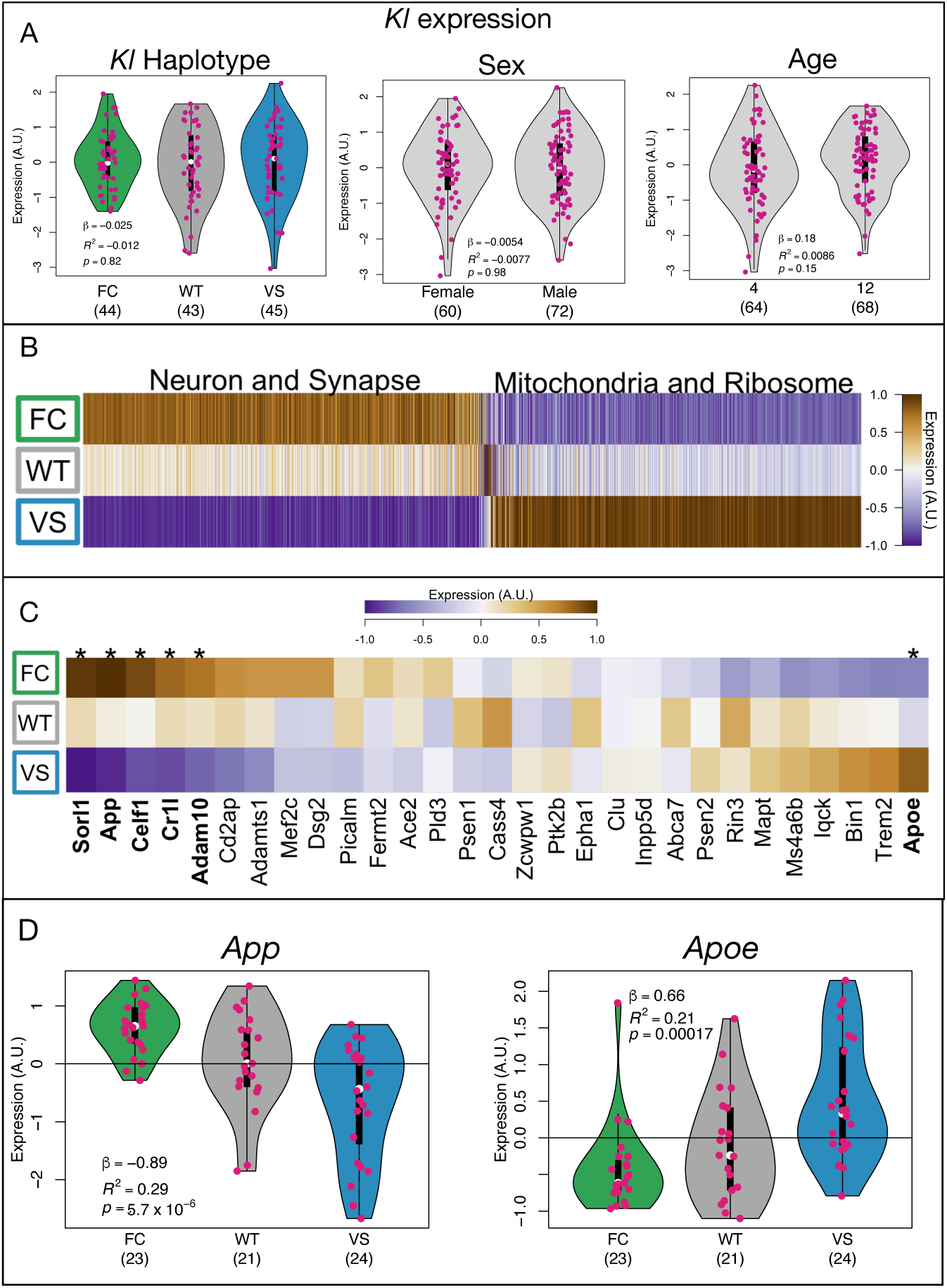
Differential gene expression related to *KL* haplotype. **A.** *KL* expression itself did not vary based on *KL* haplotype, sex or age. Numbers in parentheses indicate the number of animals in each group. **B.** There was widespread differential expression related to *KL* haplotype in the 12-month-old animals. Differentially expressed genes clustered into two general clusters. Cluster 1 was relatively down-regulated in the VS carriers and was enriched for genes involved in neuronal and synaptic functions. Cluster 2 was relatively up-regulated in VS carriers and was enriched for genes involved in mitochondrial and ribosomal functions. **C** Mean expression of genes with well-established links to AD across *KL* haplotypes. Those with significant differential expression (FDR ≤ 0.05) are marked with an asterisk and their names are bolded. **D** Detail of *App* and *Apoe* expression in the 12-month-old animals across the *KL* haplotypes. Numbers in parentheses indicate the number of animals in each group.

### *KL* haplotype had widespread effects on gene expression in the aging brain

Although *KL* expression itself was not altered in the brain, the humanized *KL* haplotypes were associated with widespread transcriptomic effects overall, suggesting altered protein function related to the amino acid changes. Table 2 shows counts of significantly differentially expressed genes (DEG) (FDR ≤ 0.05) for different factors in different age groups.

**Table 2:** Differentially expressed gene counts. A colon separating the names of factors indicates an interaction term between two factors. A dash indicates where a term was not present in the model.

| Cohort | sex | Kl haplotype | sex:Kl haplotype | age | sex:age | age:Kl haplotype |
| --- | --- | --- | --- | --- | --- | --- |
| 4 months | 8 | 9 | 0 | — | — | — |
| 12 months | 278 | 4177 | 0 | — | — | — |
| All | 65 | 4584 | 0 | 958 | 47 | 2 |

More genes had differential expression related to *KL* haplotype than had differential expression related to either sex or age. This effect of *KL* haplotype on transcription was seen almost entirely in the older age group (4177 DEG in 12 months vs. 9 DEG at four months) (Table 2). We therefore focused on the 12-month cohort for the remainder of this analysis.

In addition to focusing on 12-month-old animals, we grouped the heterozygous and homozygous carriers of each humanized allele together (Supplemental Figure 2). Within each allele, the heterozygous and homozygous carriers did not differ in measured gene expression, and more variance in gene expression was explained by a dominant genotype coding than an additive coding (Supplemental Figure 2). We thus grouped the carriers of each allele together to simplify the models and to increase power to detect differences.

Because previous studies in humans have noted interactions between *KL* haplotype and sex ^42;19^, we looked specifically for interactions between *KL* haplotype and sex in brain transcriptomics in the mice. We did not observe a significant interaction effect between *KL* haplotype and sex on brain gene expression (Table 2).

### The FC and VS alleles had opposing effects on transcription relative to the mouse wild type allele

Differentially expressed genes clustered into two groups (Fig. 2B): Cluster 1 genes had relatively high expression in the animals carrying the VS allele and were enriched for mitochondrial and ribosomal functions. Cluster 2 genes had relatively low expression in animals carrying the VS allele and were enriched for neuronal and synaptic functions. (See Supplemental Fig. 3 for more detailed reporting of functional enrichments and File1_Fig2B_Data.txt for a list of genes in each cluster).

Among the differentially expressed genes were multiple genes that have well established relationships to AD pathology ^43^ including *App*, *Apoe*, *Sorl1*, *Celf1*, *Cr1l*, and *Adam10* (Fig. 2C).

### *KL* haplotype influenced gene expression in multiple AD-related Biodomains

To further characterize the transcriptional effects of the *KL* haplotypes in relation to AD, we investigated whether the humanized *KL* haplotypes caused differential expression in Biodomains and subdomains associated with AD (Methods).

To assess the strength of the effect of the *KL* haplotypes on each biological subdomain, we fit a linear model that explained mean domain gene expression using *KL* haplotype as an ordered categorical variable (Methods). We used the variance explained by each model as a measure of the strength of the effect of the *KL* haplotypes on that subdomain. Overall, the *KL* haplotypes were most strongly associated with mean expression in the RNA spliceosome, mitochondrial metabolism and oxidative stress domains, although there was wide variation across subdomains within each domain (Fig. 3A). The VS carriers tended to have higher expression than the FC carriers of the RNA Spliceosome (linear regression: 𝑛_𝐹𝐶_ = 23, 𝑛_𝑊𝑇_ = 21, 𝑛_𝑉 𝑆_ = 24, 𝛽 = 0.14, 𝑅^2^ = 0.17, 𝑝 = 2.8 × 10^−4^) and Mitochondrial Metabolism (linear regression: 𝑛_𝐹𝐶_ = 23, 𝑛_𝑊𝑇_ = 21, 𝑛_𝑉 𝑆_ = 24, 𝛽 = 0.15, 𝑅^2^ = 0.15, 𝑝 = 4.7 × 10^−4^) domains (Fig. 3B), but lower expression in the Synapse domain (linear regression: 𝑛_𝐹𝐶_ = 23, 𝑛_𝑊𝑇_ = 21, 𝑛_𝑉 𝑆_ = 24, 𝛽 = -0.081, 𝑅^2^ = 0.14, 𝑝 = 5.5 × 10^−4^).

**Figure 3:**
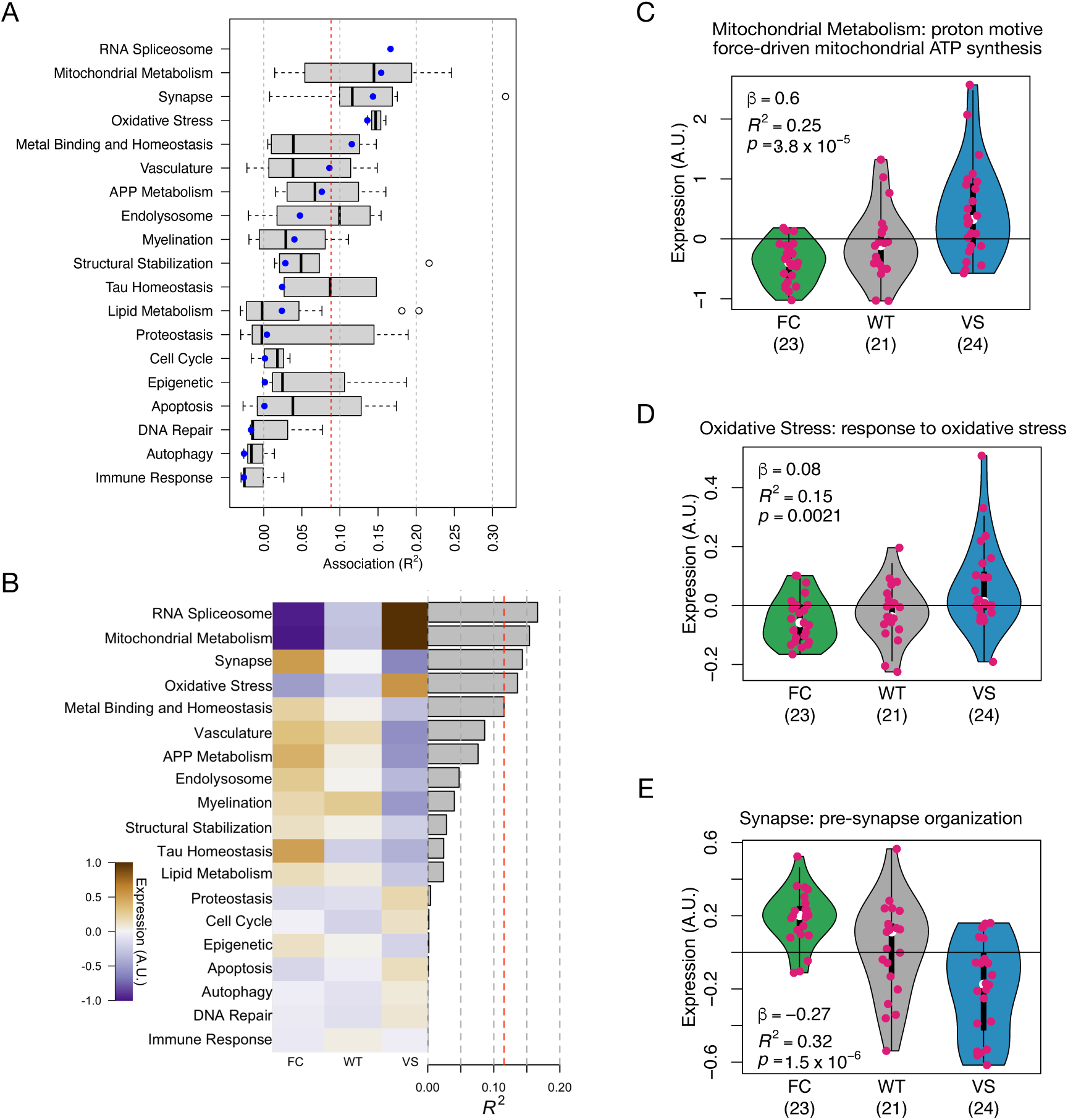
Genes grouped by AD-related Biodomains were differentially expressed across KL haplotypes. **A.** The distribution of associations between subdomains within a given Biodomain and *KL* haplotype (see Methods). The blue dot shows the association between the Biodomain as a whole and *KL* haplotype. RNA spliceosome did not have any subdomains. The vertical red line shows the FDR threshold of 0.05 **B.** Overview showing average expression of each Biodomain for each *KL* haplotype group. Gray bars show the 𝑅^2^ from a linear model testing the relationship between allele carrier status and mean gene expression. Gray vertical lines show increments of 0.05. The red line shows an FDR threshold of 0.05. **C-E.** Violin plots show expression values of genes within a subdomain. Each dot represents a single mouse, and shows the average gene expression across all genes in the subdomain. The color of each violin indicates the *KL* haplotype of the mouse. The strength of the association between subdomain gene expression and *KL* haplotype is given as 𝑅^2^ with a 𝑝-value. The three subdomains represented are the most strongly associated subdomain from Mitochondrial Metabolism (C) Oxidative Stress (D), as well as the top overall subdomain, which was presynapse organization in the Synapse Biodomain (E).

The most strongly *KL*-associated subdomain in the mitochondrial metabolism domain was “proton motive force-driven mitochondrial ATP synthesis” (Fig. 3C). Gene expression of this subdomain was significantly higher in the VS carriers than the FC carriers (linear regression: 𝑛_𝐹𝐶_ = 23, 𝑛_𝑊𝑇_ = 21, 𝑛_𝑉 𝑆_ = 24, 𝛽 = 0.6, 𝑅^2^ = 0.25, 𝑝 = 9.3 × 10^−6^). The most strongly associated subdomain in the oxidative stress domain was “response to oxidative stress” (linear regression: 𝑛_𝐹𝐶_ = 23, 𝑛_𝑊𝑇_ = 21, 𝑛_𝑉 𝑆_ = 24, 𝛽 = 0.08, 𝑅^2^ = 0.15, 𝑝 = 9 × 10^−4^) (Fig. 3D). Gene expression of this subdomain was similarly more highly expressed in VS carriers.

The subdomain that was most highly associated with *KL* haplotype was “presynapse organization” in the Synapse domain (linear regression: 𝑛_𝐹𝐶_ = 23, 𝑛_𝑊𝑇_ = 21, 𝑛_𝑉 𝑆_ = 24, 𝛽 = -0.27, 𝑅^2^ = 0.32, 𝑝 = 2.7 × 10^−7^) (Fig. 3E). For this subdomain, the highest expression was in the FC carriers.

### *KL* haplotypes caused differential splicing of genes in glutamatergic synapses

Because the RNA spliceosome Biodomain was the most highly differentially expressed across *KL* haplotypes, we investigated differential splicing between 12-month-old mice carrying the FC and VS haplotypes (Methods). We identified 1831 genes with differential exon usage at an FDR of 0.05. This group of genes was highly enriched for GO terms related to synapse, and glutamatergic synapse in particular (Supplemental Figure 4). Other processes that were enriched in the differentially expressed genes, such as mitochondrial metabolism and ribosomal genes were absent from this group of differentially spliced genes.

### Neurodegeneration and oxidative phosphorylation KEGG pathways were differentially expressed by *KL* haplotype

In addition to the AD-specific Biodomains, we investigated the effects of *KL* haplotypes on gene expression in KEGG pathways ^32;33^. KEGG pathways provide a more disease-agnostic grouping of genes in a way that also provides a high level of molecular detail about the relationships between genes, and their involvement in specific signaling pathways.

We assessed the relationship between *KL* haplotypes and KEGG pathways using the method described above for Biodomains. At an FDR of 0.05, 153 of the total 356 KEGG pathways were significantly differentially expressed. These included multiple neurodegenerative pathways including those labeled as relevant to prion disease, Parkinson’s disease, ALS, and Huntington’s disease (Fig. 4A). KEGG pathways related to glutamatergic synapses and oxydative phosphorylation mirrored mitochondrial metabolism (Fig. 4C) and synaptic (Fig. 4E) differential expression seen in the Biodomain analysis.

**Figure 4:**
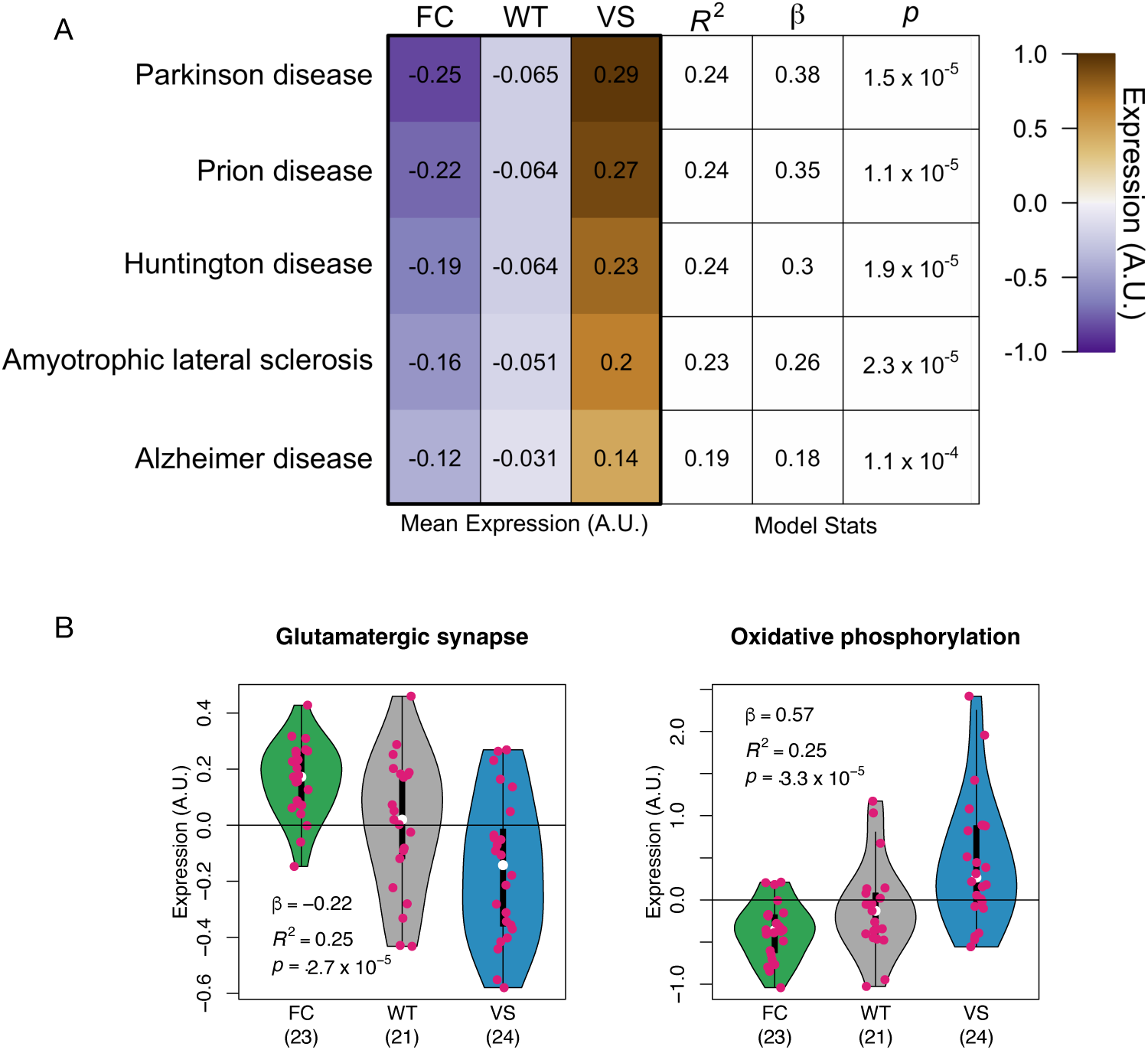
Expression variation in KEGG pathways. **A** Results for KEGG neurodegeneration pathways. Left three columns show the mean expression of the genes in each KEGG pathway for each *KL* haplotype. The right three columns show the stats for the linear models explaining gene expression with *KL* haplotype. **B** Genes related to oxidative phosphorylation were more highly expressed in VS carriers relative to FC carriers. Genes related to glutamatergic synapses had reduced expression relative to FC carriers. Both results mirror those seen in the Biodomains.

### Intersectional analysis identifies genes related to APP Metabolism were differentially expressed across *KL* haplotypes

To combine the AD specificity of Biodomains with the biochemical annotations in KEGG pathways, we considered the intersecting set of genes between each Biodomain and KEGG pathway. These intersections provided both molecular interaction information as well as AD relevance and have been shown in a more general context to improve pathway analyses of transcriptomic data ^44^.

One example of enhanced specificity is refining the association between *KL* haplotypes and MAPK signaling to its known role in apoptosis ^45^ (Fig. 5). The 133 genes that intersect the MAPK signaling pathway (map04010) and the apoptosis Biodomain were differentially expressed across *KL* haplotypes (linear regression: 𝑛_𝐹𝐶_ = 23, 𝑛_𝑊𝑇_ = 21, 𝑛_𝑉 𝑆_ = 24, 𝛽 = -0.11, 𝑅^2^ = 0.15, 𝑝 = 4.2 × 10^−4^) (Fig. 5A left), whereas the 27 genes common for MAPK signaling and the DNA repair Biodomain were not (linear regression: 𝑛_𝐹𝐶_ = 23, 𝑛_𝑊𝑇_ = 21, 𝑛_𝑉 𝑆_ = 24, 𝛽 = -0.014, 𝑅^2^ = -0.028, 𝑝 = 0.71) (Fig. 5A right). This suggests that this level of detail is meaningful in this context and can narrow the differential gene expression analysis beyond that of the broad KEGG pathways and Biodomains. More specifically, this result suggests that the known influence of *Kl* on MAPK-mediated apoptosis ^45^ is altered by the common *KL* haplotypes.

**Figure 5:**
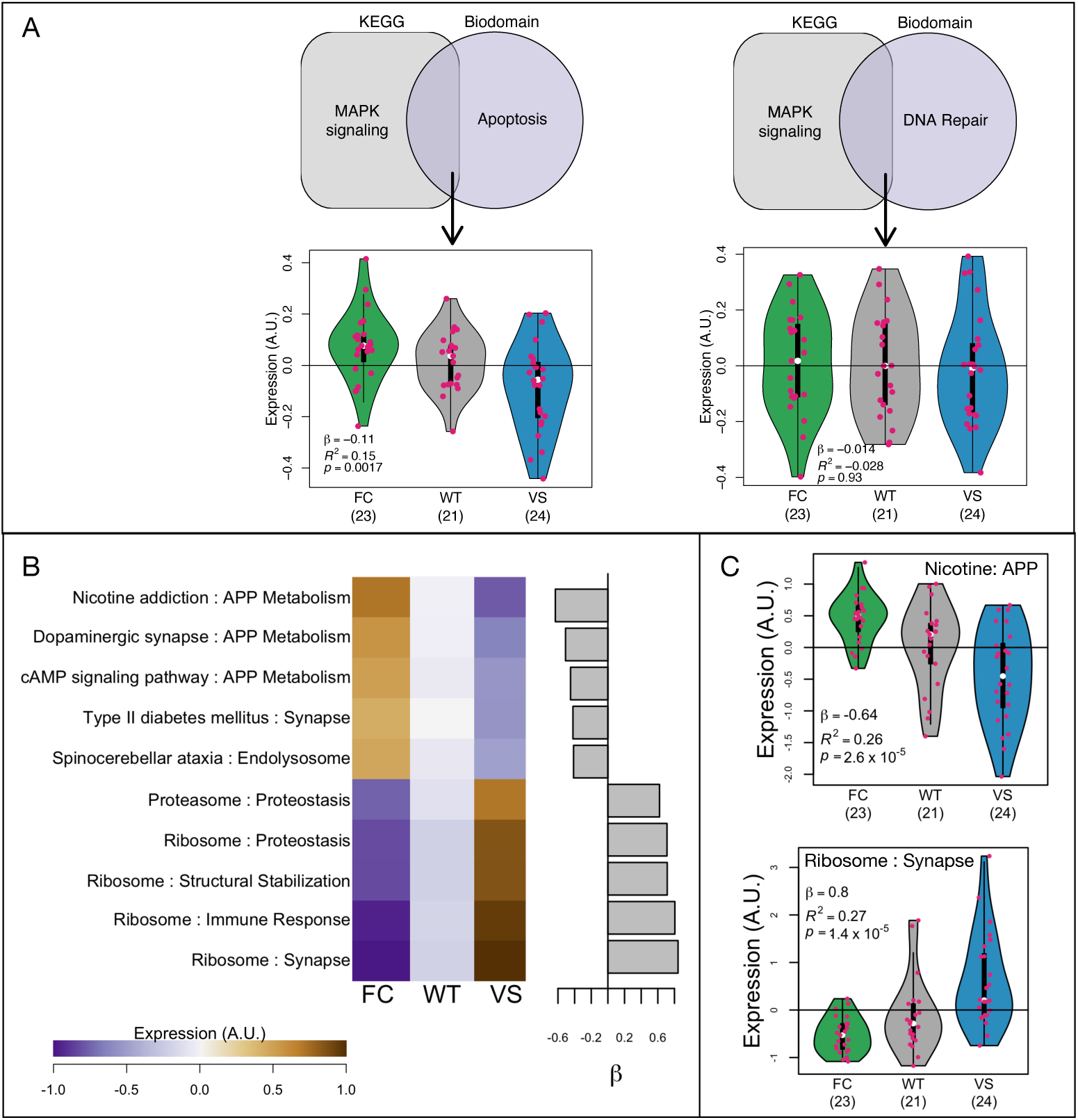
Intersections of Biodomains and KEGG pathways provide additional information. **A** Shaded figures show intersections between specific KEGG pathways and Biodomains. (left) Genes in the intersection of the KEGG pathway MAPK signaling and the Biodomain Apoptosis are differentially expressed across the *KL* haplotypes (linear regression: 𝑛_𝐹𝐶_ = 23, 𝑛_𝑊𝑇_ = 21, 𝑛_𝑉 𝑆_ = 24, 𝛽 = -0.11, 𝑅^2^ = 0.15, 𝑝 = 4.2 × 10^−4^). (right) Genes in the intersection of the KEGG pathway MAPK signaling and the Biodomain DNA repair are not differentially expressed (linear regression: 𝑛_𝐹𝐶_ = 23, 𝑛_𝑊𝑇_ = 21, 𝑛_𝑉 𝑆_ = 24, 𝛽 = -0.014, 𝑅^2^ = -0.028, 𝑝 = 0.71). **B.** Mean expression of the top significant pathway intersections ordered by *KL* haplotype effect size. Brown and purple colors indicate increased and decreased expression respectively. The bar plot shows the *KL* haplotype effect size on each pathway intersection. Intersections with positive effects are more highly expressed in VS carriers. Those with negative effects have lower expression in VS carriers. **C.** Examples of intersections with positive and negative effect sizes to show differential expression in more detail.

For each KEGG-Biodomain intersection, we calculated mean expression across all individuals and used a linear model as described above to assess the effect of haplotype on the expression of each intersectional gene set. At an FDR of 0.001 there were 180 pathway intersections associated with *KL* haplotype. Positive effects indicated that the highest gene expression was in the VS carriers and the lowest was in the FC carriers (Figure 5B). Negative effects indicated the reverse, that the lowest expression was in the VS carriers and the highest was in the FC carriers (Figure 5B). Gene lists and the statistical results for each intersection are provided in File2_Intersection_Stats.txt.

Clustering the significant intersections by effect highlighted a number of patterns of interest (Figure 6). KEGG pathways were affected by *KL* haplotype consistently across multiple intersections with Biodomains, suggesting a high degree of functional coherence in each. The Ribosome KEGG pathway, for example, had strong positive coefiicients across multiple Biodomains. In contrast, Biodomains had both positive and negative responses to *KL* haplotype across intersections. For example, Mitochondrial Metabolism genes related to ribosome and oxidative phosphorylation had higher expression in the VS animals, whereas Mitochondrial Metabolism genes related to synapses and hormonal signaling had lower expression in the same animals. This result suggests that rather than being up- or down-regulated overall, subsets of mitochondrial genes were either up- or down-regulated in a manner related to specific biological processes. Of particular interest in the context of AD, were a group of neurodegenerative KEGG pathways intersecting the Mitochondrial Metabolism Biodomain that all had increased expression in VS carriers (highlight Fig. 6). This result suggests that two common human *KL* haplotypes potentially have an effect on mitochondrial function directly related to neurodegenerative disease ^46–50^. This is granularity of analysis was not apparent from using the biological subdomains alone (Supplemental Figure 5)

**Figure 6:**
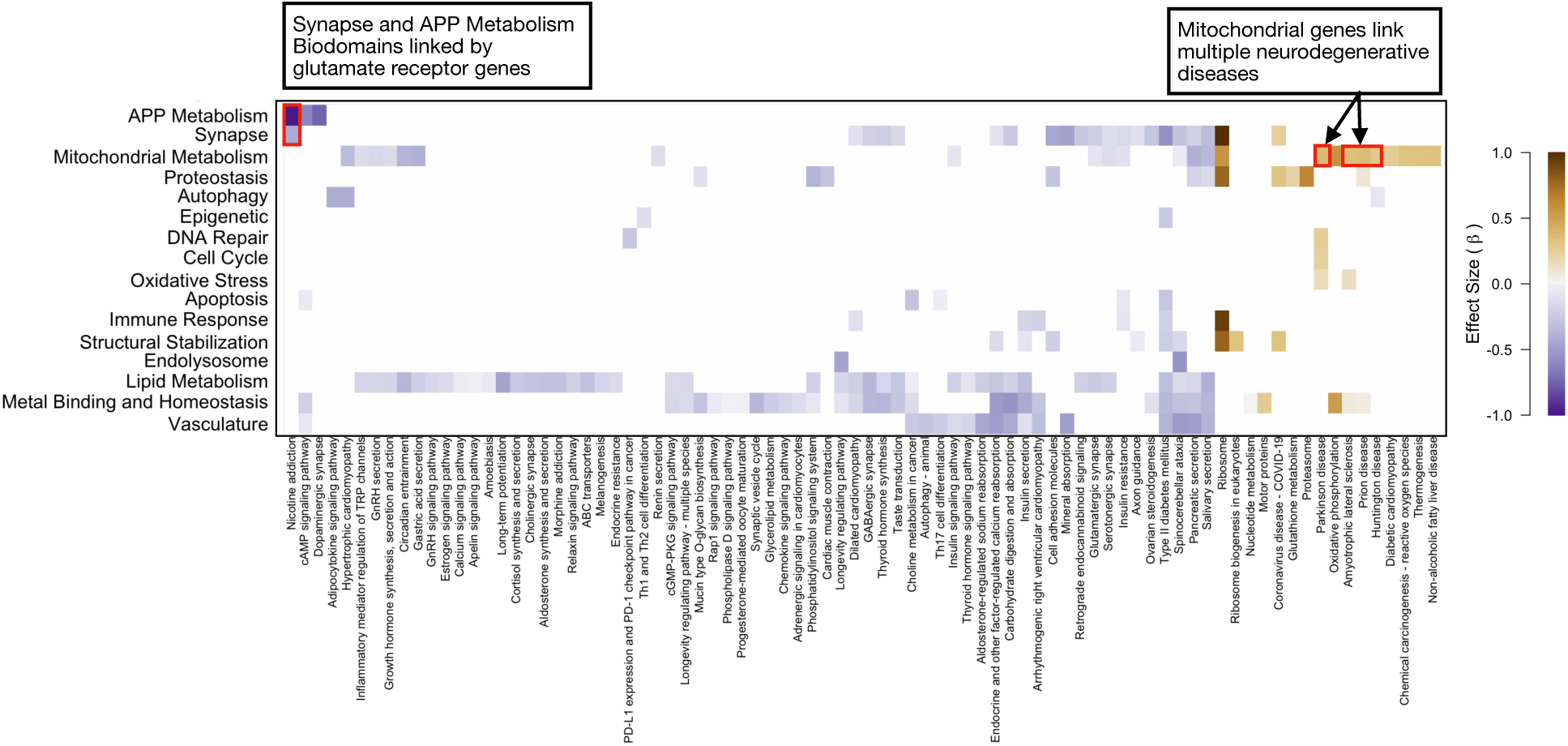
Biadjacency matrix of significantly differentially expressed KEGG-Biodomain intersections at FDR ≤ 0.001. Cell colors indicate effect direction and magnitude of haplotype on the mean gene expression of the group. Positive values indicate higher expression in VS animals and negative values indicate lower expression in VS animals. Red boxes with related text highlight intersections of interest.

The intersections for which haplotype had the strongest effect were between the Biodomain APP Metabolism and the KEGG pathway Nicotine addiction (red box in Figure 6). The term Nicotine addiction also had a significant intersection with the Synapse Biodomain thereby functionally connecting APP Metabolism with synaptic function. The intersection of these three terms contained 10 genes highly enriched for glutamate receptor components including *Grin2b* (Figure 7A). *Grin2b* (also called *GluN2B*) encodes a subunit of the NMDA receptor (NMDAR), which plays a key role in learning and memory ^51^ and mediates the effects of *Kl* on depressive symptoms in mice ^52^. KL increases *Grin2b* expression in synapses ^4^ while simultaneously protecting neurons from glutamate excitotoxicity ^53^. KL also reduces the depletion of NMDAR otherwise observed in humanized APP mice ^54^ thereby directly connecting *Kl* and *Grin2b* to APP metabolism.

**Figure 7:**
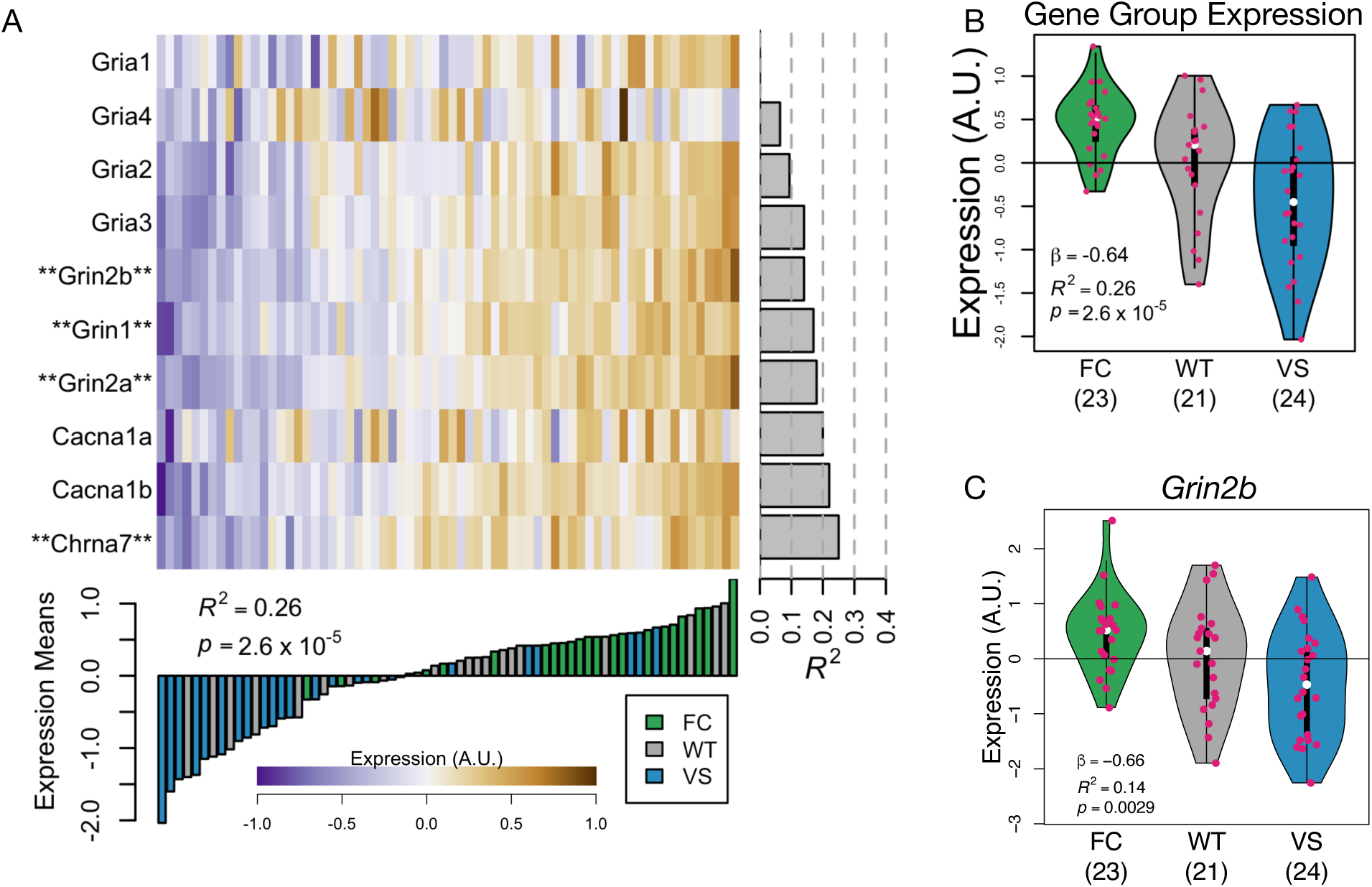
Genes in the intersection of the Biodomains Synapse and APP Metabolism and the KEGG pathway Nicotine addiction. **A.** Heat map shows expression of individual genes in the intersection of the Biodomains Synapse and APP Metabolism and the KEGG pathway Nicotine addiction. Gene names marked with asterisks are in the KEGG pathway Alzheimer disease (map05010). The bar plot below the heat map shows the mean expression of the group of genes overall for each individual animal. The bar color indicates the animal’s haplotype. The gray bars on the right-hand side of the heat map show how much variance in each gene’s expression is explained by *KL* haplotype. **B.** Mean expression of the group of genes is highest in FC carriers and lowest in VS carriers. **C.** *Grin2b*, a mediator of *Kl*’s effects in the brain was differentially expressed across haplotypes.

Notably, although the RNA spliceosome Biodomain had the strongest association with *KL* haplotype overall, this domain did not appear in this intersectional analysis. The top two KEGG intersections for the RNA Spliceosome domain were RNA degradation (linear regression: 𝑛_𝐹𝐶_ = 23, 𝑛_𝑊𝑇_ = 21, 𝑛_𝑉 𝑆_ = 24, 𝛽 = 0.44, 𝑅^2^ = 0.22, 𝑝 = 4 × 10^−5^), and Spliceosome (linear regression: 𝑛_𝐹𝐶_ = 23, 𝑛_𝑊𝑇_ = 21, 𝑛_𝑉 𝑆_ = 24, 𝛽 = 0.31, 𝑅^2^ = 0.21, 𝑝 = 4.1 × 10^−5^), whose 𝑝 values were slightly above the stringent significance threshold we used for the intersections. Both sets of genes were more highly expressed in the VS animals (Supplemental Figure 6). This result highlights the importance of the different perspective provided by the intersectional analysis. Although spliceosome-related transcripts were differentially expressed across the haplotypes, there were additional important processes identified by the intersectional analysis as being even more highly differentially expressed that would not have been identified without this functional granularity.

### Transcriptomic effects in mice are consistent with results in human brain organoids

To investigate the translatability of these results to human brains, we used transcriptomic data collected in human brain organoids overexpressing *KL* ^28^. We calculated the transcriptomic effects of *KL* overexpression on the KEGG-Biodomain intersections and compared these to the transcriptomic effects of *Kl* haplotype we observed in mice (Methods). The strongest effects of *Kl* haplotype in mice were recapitulated in the human brain organoids (Fig. 8A). In mice, animals with the VS haplotype had reduced expression of glutamate receptor genes (APP Metabolism - Nicotine addiction intersection), and increased expression of ribosomal genes in the Synapse Biodomain (Fig. 5C). In the human brain organoids, overexpression of *KL* had the same effects on these two groups of genes (Fig. 8A), suggesting that the VS allele increased KL protein abundance in the mouse brains.

**Figure 8:**
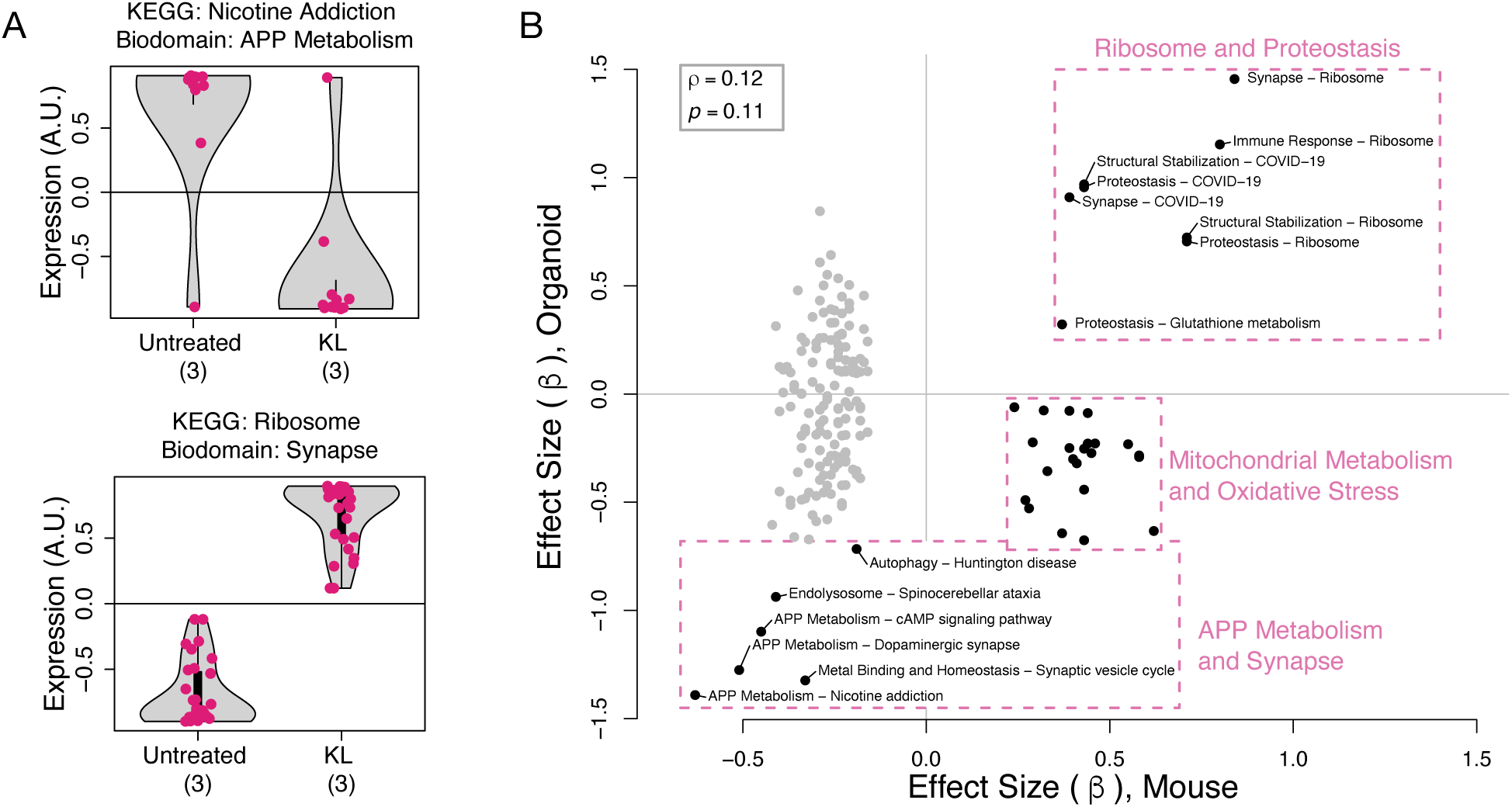
Comparison of effects of the VS allele in mice and *KL* induction in human iPSC-derived brain organoids. **A.** Induction of *KL* expression in 13-week-old human brain organoids caused a reduction in mean expression of glutamate receptor genes (top) and an increase in mean expression of ribosomal genes involved in synaptic processes (bottom). Compare to Fig. 5C. Each dot represents the mean expression of a single gene across three organoid replicates. **B.** Comparison of *KL* effects on gene expression in mouse brain and brain organoids across KEGG-Biodomain intersections. Each dot compares the 𝛽 coefiicients from linear models describing the effect of either the VS allele (𝑥-axis) or *KL* induction in human brain organoids (𝑦-axis) on mean expression of the group of genes defined by the KEGG-Biodomain pathway intersection. The overall correlation between the effect sizes of the VS allele or *KL* induction was not significant (Spearman 𝜌 = 0.12, 𝑝 = 0.11). All values are available in File3_Organoid_Comparison.txt.

Across all KEGG-Biodomain intersections there was no significant correlation between the mouse and organoid transcriptional variation (Fig. 8B) (Spearman 𝜌 = 0.12, 𝑝 = 0.11). However, multiple KEGG-Biodomain intersections containing genes related to APP Metabolism and Synapse, as well as those related to Ribosomes and Proteostasis were strongly similar in the mouse and human model systems (Fig. 8B boxes). One notable difference between the mouse and organoid responses was in groups of genes related to mitochondrial metabolism and oxidative stress (Fig. 8B). Many of these groups of genes had reduced expression in response to *KL* overexpression in organoids, but had increased expression in the VS mice relative to the FC mice. Such differences could be attributable to many factors. For example, the nature of the experimental intervention is different: increased *KL* expression compared to *KL* haplotype variation. Although the VS allele may increase KL abundance, it may alter the processing or the function of the protein as well. The responses of the two systems may be cell-type specific and the mouse brains contain cell types not present in the organoids. The organoids, moreover do not interact with the other organ systems, which themselves respond to KL and produce factors that influence the brain ^55^. This organoid study also did not look specifically at the effects of oxidative stress, which must be induced in organoids ^56^ in contrast to aging organisms, in which it occurs naturally. Finally, there could be species-specific differences in response to

*KL*. These experimental differences may also explain the cluster of gray points in Fig. 8B that were enriched in lipid metabolism and insulin signaling genes and had discordant responses in the mouse and organoid experiments. Given the many differences between the systems, it is remarkable that the glutamate receptor genes and the ribosomal genes had comparable responses in the two models. This may indicate that these processes are neuron-specific effects of *KL* that can be captured by the organoid model.

## Discussion

Here we demonstrated that humanized FC and VS haplotypes of the gene Klotho (*KL*) introduced to C57BL/6J mice differentially affected the brain transcriptome including many pathways associated with neurodegenerative disorders. This work supports the hypothesis that these haplotypes alter human aging and the risk of developing age-related neurodegenerative disorders of the brain. There are multiple possible mechanisms for the observed effects of the *KL* haplotypes. For example, the haplotypes may cause variation KL protein secretion. The concordance between the transcriptomic variation in the mouse brains and transcriptomic variation in human brain organoids with induced *KL* overexpression suggests that at least part of the observed variation in mice may be due to neuron-intrinsic excess abundance of KL protein. Alternatively, the variants may cause variation in the relative abundance of KL protein isoforms. Future work will examine the abundance of KL isoforms in the humanized *KL* mice to further investigate the mechanisms underlying the haplotype effects observed here.

Multiple genes with strong experimental ties to AD, including *App*, *Apoe*, and *Sorl1*, were differentially expressed across the *KL* haplotypes and multiple others (e.g. *Trem2*, *Cd2ap*, *Adamts1*, and *Bin1*) exhibited changes that did not reach the genome-wide false discovery rate (Figure 2). That key AD genes were differentially expressed in the presence of naturally occurring *KL* variants, as opposed to an engineered null or overexpression *Kl* mutations, is of potential translational relevance to AD risk modification. In particular, the VS allele has been implicated in reduced risk of AD in *APOE4* carriers ^15^ but the association is still unresolved ^20^. These findings motivate further studies of the interaction between these *KL* alleles and humanized *APOE3* and *APOE4* alleles in mice, where we hypothesize that the presence of *APOE3* will attenuate the *KL* haplotype effects ^57;15^.

Over the entire transcriptome, we observed major changes in genes influencing mitochondrial, ribosomal, and synaptic pathways, each of which is critical to cognitive function and impaired in AD ^47^. For example, mitochondrial fission and fusion, which are critical for maintenance of healthy mitochondrial function, are disrupted in AD leading to altered morphology and distribution of mitochondria in the neurons of AD-affected brains ^49;58^. The degradation of mitochondrial function impairs neurotransmitter synthesis ^59;50^, calcium concentration and signal transduction ^60;61^ and increases oxidative stress ^48^, all of which are hallmarks of AD ^49;47;46^. Ribosomal impairment is seen early in AD during the mild cognitive impairment (MCI) phase ^62^. Impairments include reduced capacity for protein synthesis, reduced rRNA and tRNA levels and increased oxidation ^62^. Furthermore, the age-related reduction of ribosomal quantity impairs proteostasis, causing an accumulation of misfolded proteins that form neurotoxic aggregates ^63–65^. To confirm alterations of ribosomal and mitochondrial processes, further histological and functional analyses of these mice are needed. Future work will examine mitochondrial size, morphology, and distribution, as well as oxidative stress levels in the *KL* mice. Ribosomal function and distribution can also be assayed to identify variation in protein synthesis or localization across the *KL* haplotypes.

That both ribosomal and mitochondrial dysfunction can cause synaptic dysfunction raises the possibility that the transcriptomic alterations we observed here in mitochondria and ribosomes cause the variation in synaptic transcripts. Alternatively, the *KL* haplotypes may independently influence synaptic density and function ^54;66^. That induction of *KL* expression in human brain organoids had the same effect on glutamate receptor genes, but the opposite effect on mitochondrial genes suggests that the synaptic effects may be independent of the mitochondrial effects. Transcriptomic measurements in the mice at time points between four months of age, when no transcriptional differences were observed, and 12 months of age, when transcripts in all three systems were altered, could potentially put in chronological order the effects of the *KL* haplotypes to better identify causal influences among the processes.

Alternatively, it is possible that observed variation in ribosomal, mitochondrial, and synaptic programs across the *KL* haplotypes indicate age-related variation in cell type composition or aging rates. Mitochondrial ^67^, and ribosomal ^68^ number and activity vary widely across cell types in the brain, and aging has been shown to alter cell type composition in mouse brains ^69^. Different cell types have furthermore been shown to be differentially susceptible to aging ^70;71^, and variation in transcription may indicate accelerated or delayed aging of individual cell types. Further histological or single-nucleus sequencing could investigate the possibility of variation in cell composition across the *KL* haplotypes.

Despite broad Alzheimer’s related effects on the transcriptome, we did not observe effects of the *KL* haplotypes on immune response genes at four or 12 months of age. Immune response genes and pathways are central in the genetics of late-onset AD ^72–74^ and are commonly upregulated in the presence of neuropathological aggregates ^75–77^. In the specific context of AD, the KEGG-Biodomain intersectional analysis did detect limited *KL* haplotype effects for KEGG pathways in the Immune Response Biodomain related to insulin secretion and type II diabetes, for which the VS mice had relatively reduced expression. Further studies of mice aged to 18 months and beyond will better determine if the *KL* haplotypes significantly affect age-related immune function, and combinations of these alleles with neuropathological amyloid and/or tau drivers might further elucidate any disease-relevant contributions.

We observed a lack of interaction between the effects of *KL* haplotype and animal sex. Previous studies in humans have identified *KL* interactions with sex ^78;42^, but cohort studies in humans are confounded by cultural variation in treatment of males and females over time, and may not be able to identify true biological effects of haplotype variation, particularly in outcomes as complex as cognition. Alternatively, sex effects may be present in humans, but not in mice, or may be present in mice in more advanced ages. Neither of these cases is possible to confirm with the present data.

In conclusion, we have used novel mouse models to demonstrate that the common human *KL* haplotypes, FC and VS, significantly alter multiple transcriptomic programs relevant to Alzheimer’s disease in the brain. These findings further link variants in the aging factor *KL* with age-related neurodegenerative disease. Our results motivate additional studies of *Kl* haplotypes in combination with humanized *APOE* alleles and neuropathological drivers to further elucidate the mechanisms of risk and protection suggested by human genetic studies.

## Supporting information

File2_Intersection_Stats

File3_Organoid_Comparison

## Acknowledgements

This work was supported by a grant from the National Institutes of Health (RF1AG075701 and U54 AG054345).

We thank the Genome Technologies scientific service at The Jackson Laboratory for the RNA sequencing and the staff at the Genetic Engineering Technologies Service at the Jackson Laboratory for their contribution.

## Supplemental Figures

**Figure S1:**
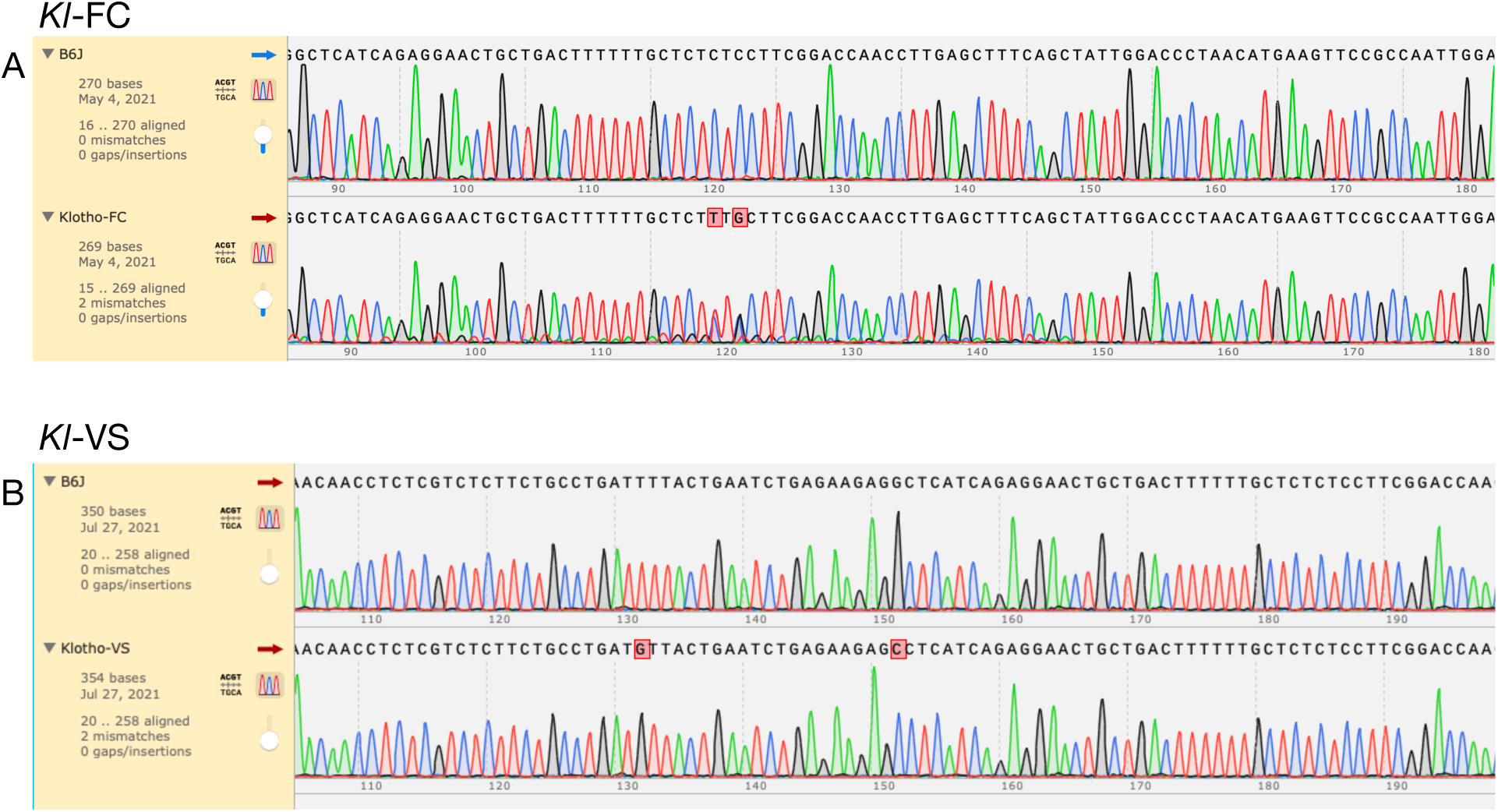
Sequencing results for humanized *Kl* alleles compared to C57BL/6J (B6) reference. **(A)** Sequenc-ing results for the *Kl*-FC construct (bottom) compared to the B6 reference (top). Highlighted bases show the SNP used to introduce the S370C variant (TCC>TGC), as well as a silent mutation (CTC>CTT) immediately downstream used to prevent re-cutting. **(B)** Sequencing results for the *Kl*-VS construct (bottom) compared to the B6 reference (top). Highlighted bases indicate the SNP used to introduce the F352V variant (TTT>GTT) as well as a silent mutation (AGG>AGC) immediately downstream used to prevent re-cutting.

**Figure S2:**
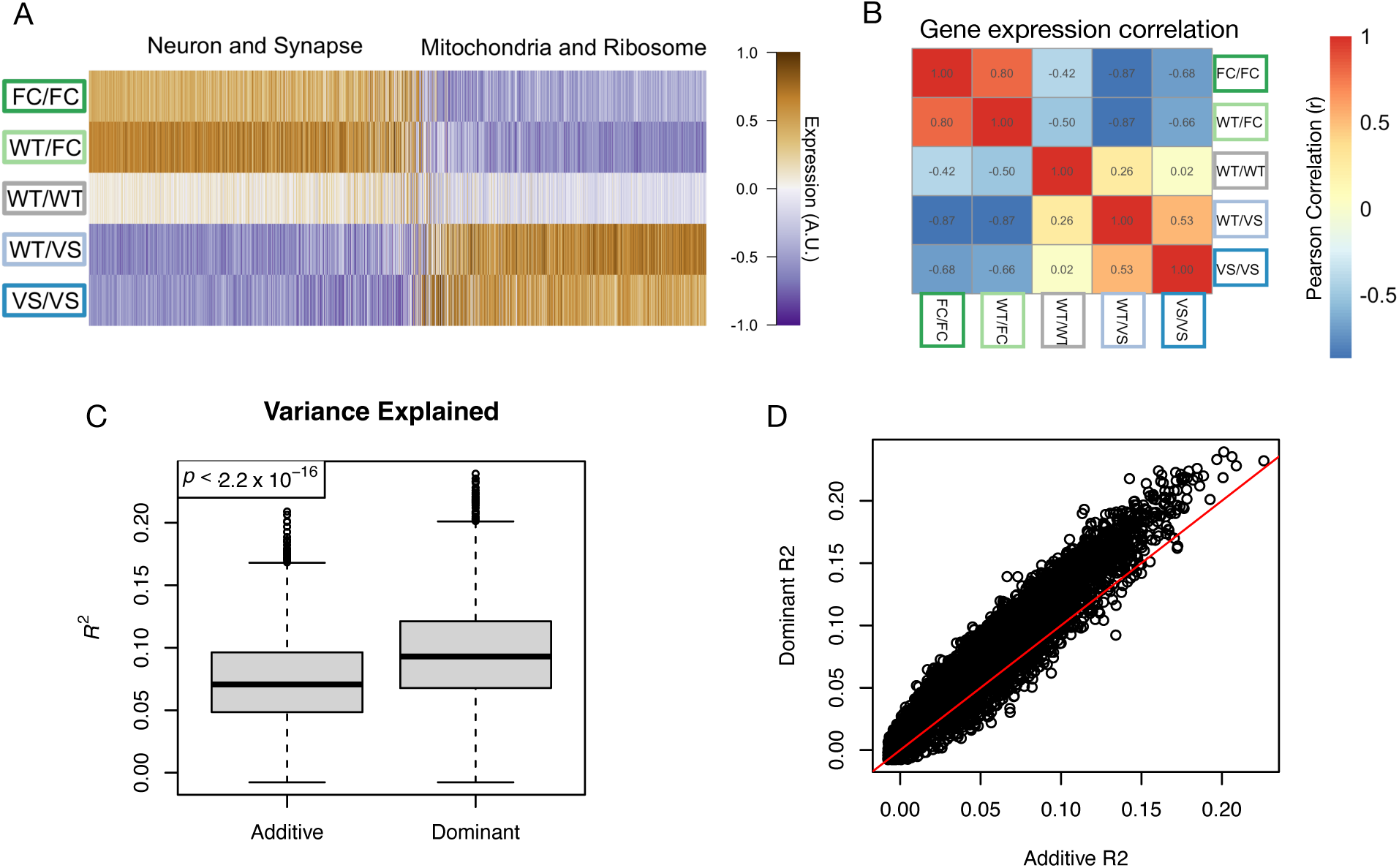
Justification for grouping allele carriers together. **A.** Mean expression for all genes significantly differentially expressed across the genotypes. Values shown are the mean expression for each genotype with genes shown in columns and genotypes shown in rows. The expression clusters into two major clusters enriched for neuronal processes and mitochondrial processes as indicated. The FC carriers tend to have correlated gene expression across the clusters, as do the VS carriers. **B.** Gene expression correlations between genotype pairs. The FC carriers are correlated with each other and the VS carriers are correlated with each other. FC and VS carriers have anti-correlated gene expression. **C.** Comparison of variance explained for additive and dominant linear models for all significantly differentially expressed genes. The dominant coding explained significnatly more variance than the additive coding. **D.** Comparison of statistics from linear models using dominant and additive codings. The dominant coding tended to have higher variance explained than the additive coding across the range of variance explained, but particularly in genes with high variance explained by genotype. Red line shows 𝑦 = 𝑥.

**Figure S3:**
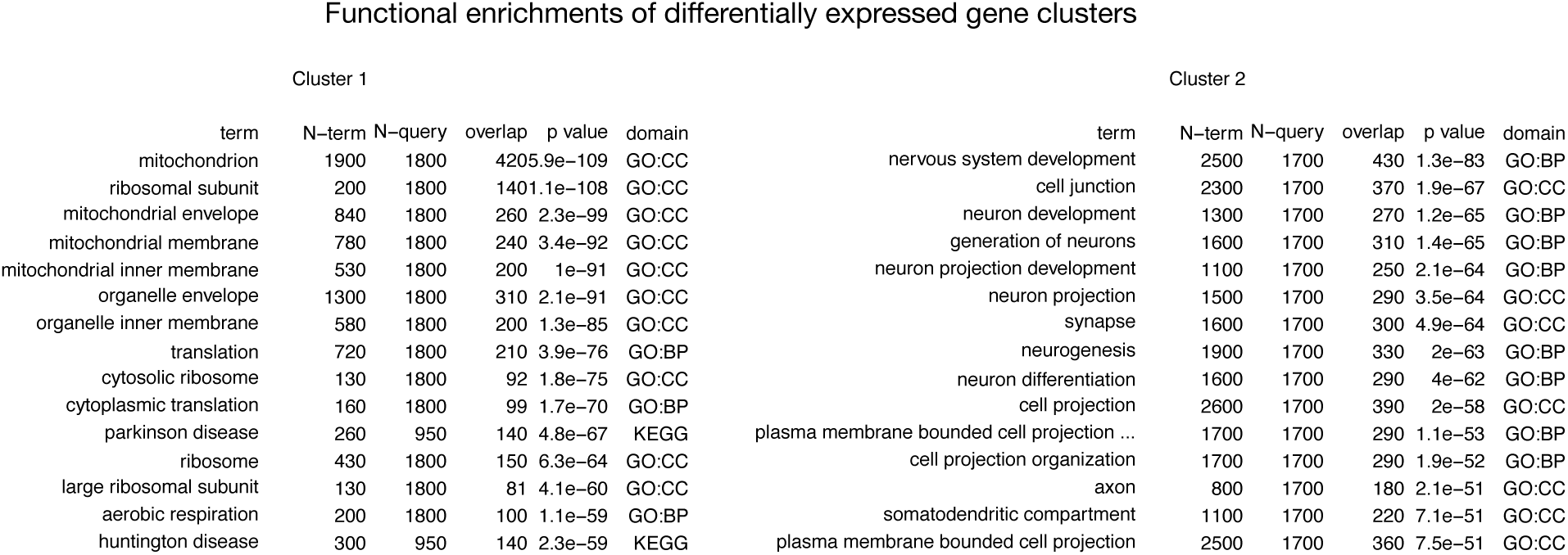
Functional enrichments of genes shown in Fig. 2B. Genes in cluster 1 were relatively up-regulated in VS carriers and were enriched in mitochondrial and ribosomal functions. The top 15 most enriched terms are shown here. Genes in cluster 2 were relatively down-regulated in VS carriers and were enriched in neuronal and synaptic functions. The top 15 most enriched terms are shown here.

**Figure S4:**
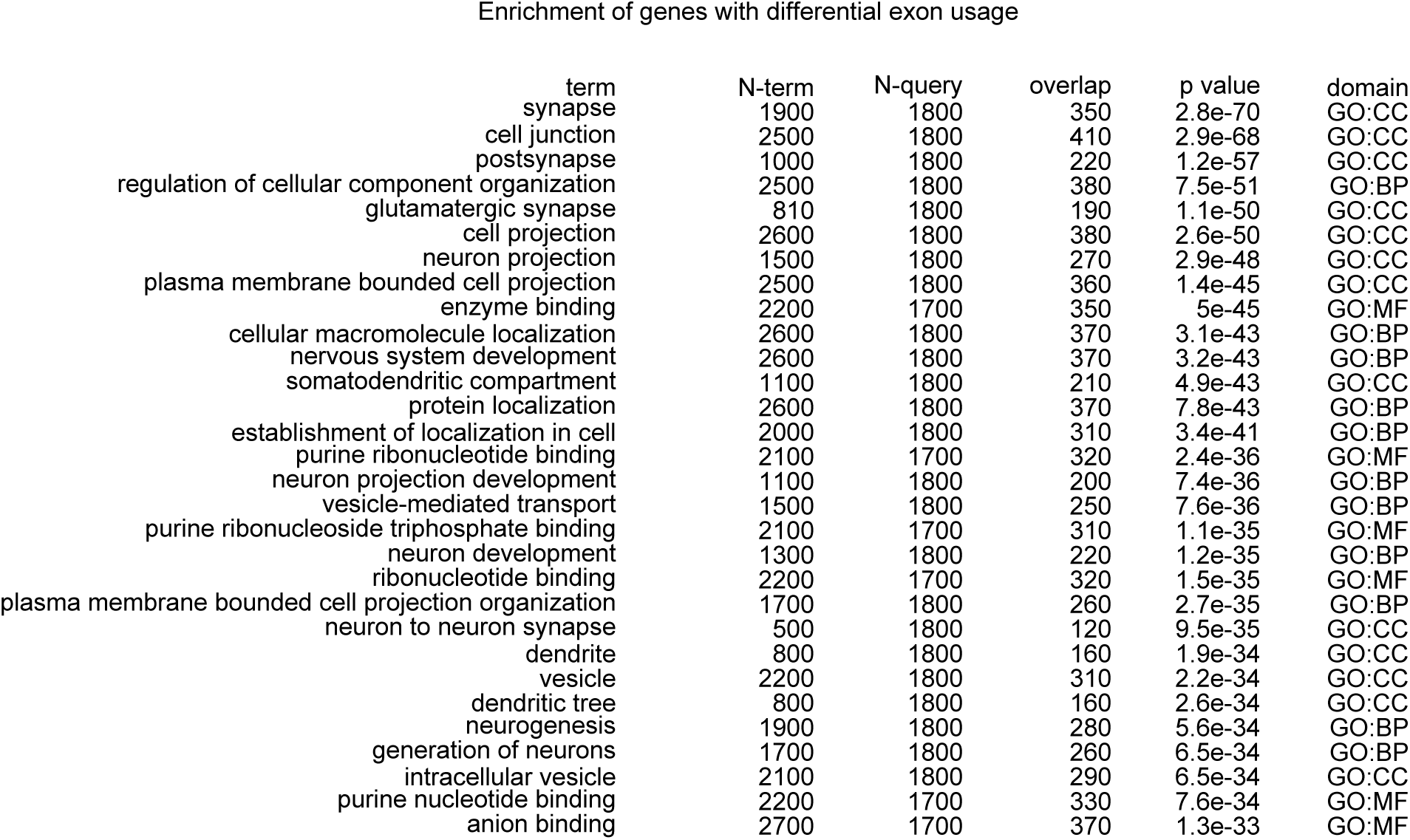
Functional enrichments of genes with significantly varying exon usage across *Kl* haplotypes.

**Figure S5:**
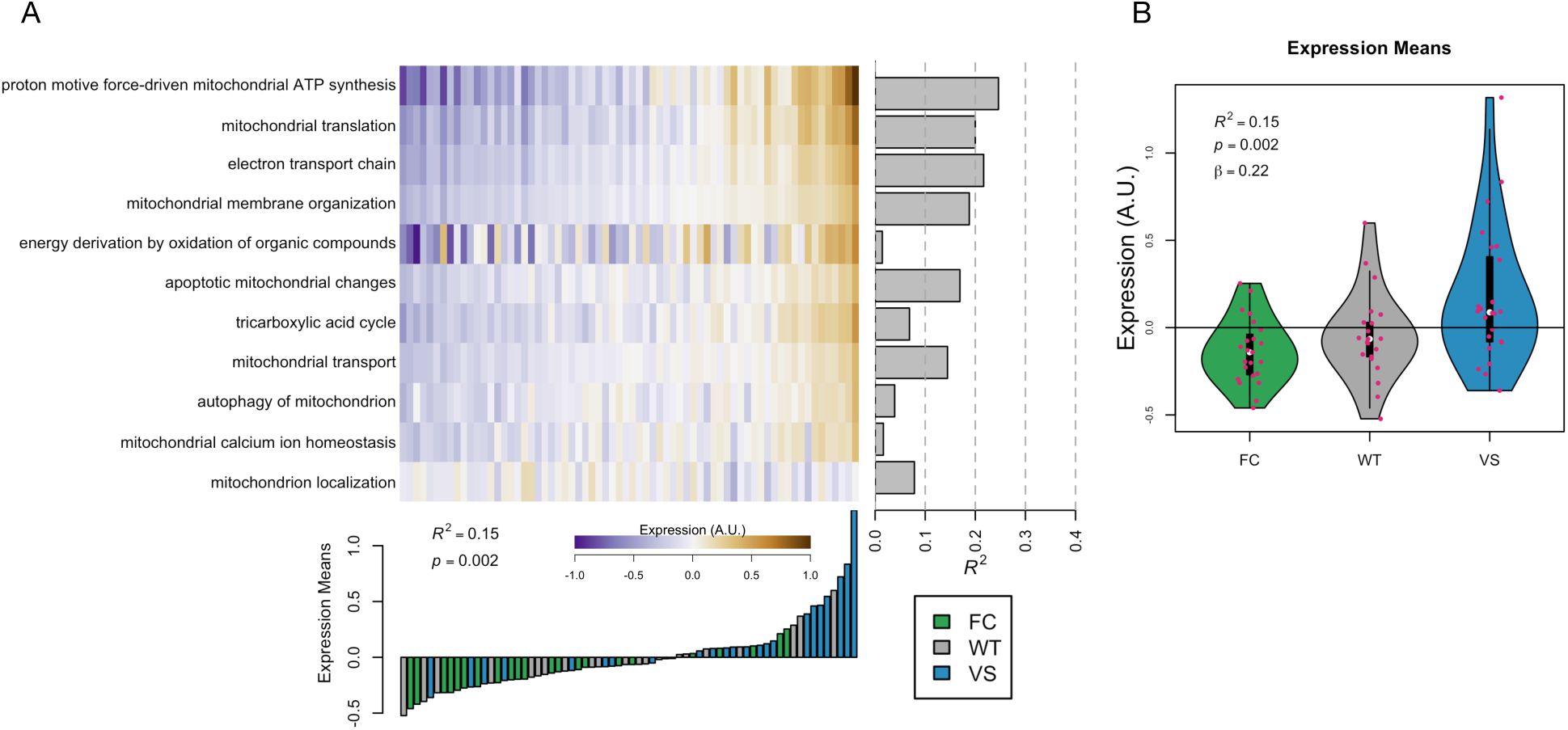
Mitochondrial metabolism subdomains. **A** (Above) Average expression of each subdomain in the Biodomain Mitochondrial Metabolism. Negative values are shown in purple, and positive values are shown in brown. Gray bars indicate the variance explained in expression by genotype. (Below) Average expression across all subdomains for each individual mouse. The color of each bar indicates the mouse’s *Kl* haplotype. **B** Violin plot showing the overall expression of the Biodomain separated by genotype.

**Figure S6:**
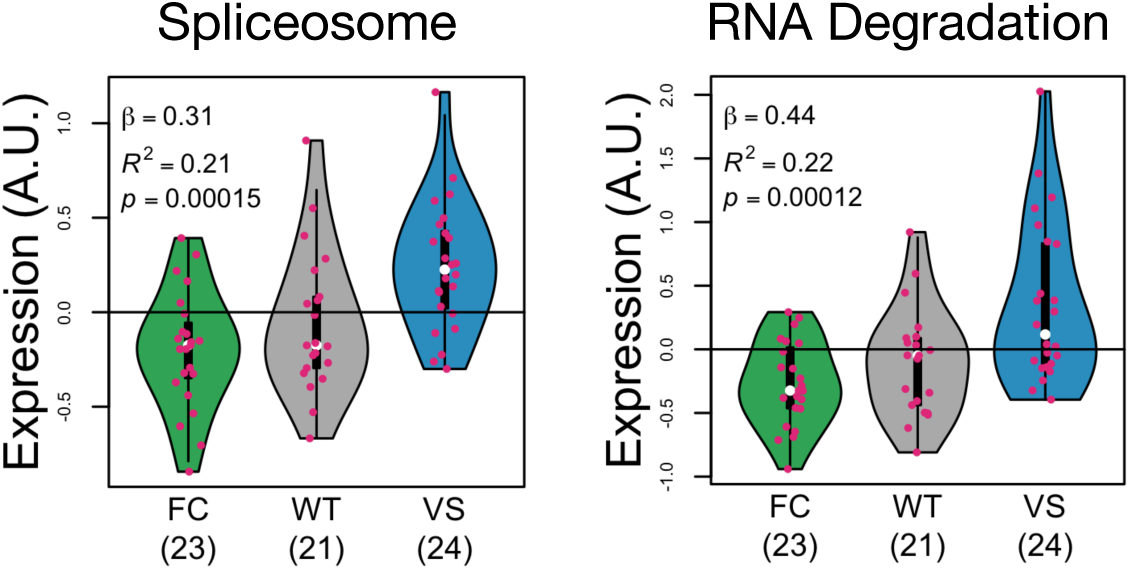
Intersections of the Biodomain RNA Spliceosome and KEGG pathways were differentially expressed, but the adjusted 𝑝 values were slightly above the cutoff used in the intersectional analysis. Titles on each panel indicate the name of the KEGG pathway that was intersected with the RNA Spliceosome Biodomain.

## References

[1] Kenneth Lim, Arnoud Groen, Guerman Molostvov, Tzongshi Lu, Kathryn S Lilley, David Snead, Sean James, Ian B Wilkinson, Stephen Ting, Li-Li Hsiao, Thomas F Hiemstra, and Daniel Zehnder. -klotho expression in human tissues. J Clin Endocrinol Metab, 100(10):E1308–18, Oct 2015. doi: 10.1210/jc.2015-1800.

[2] M Kuro-o, Y Matsumura, H Aizawa, H Kawaguchi, T Suga, T Utsugi, Y Ohyama, M Kurabayashi, T Kaname, E Kume, H Iwasaki, A Iida, T Shiraki-Iida, S Nishikawa, R Nagai, and Y I Nabeshima. Mutation of the mouse klotho gene leads to a syndrome resembling ageing. Nature, 390(6655):45–51, Nov 1997. doi: 10.1038/36285.

[3] Hiroshi Kurosu, Masaya Yamamoto, Jeremy D Clark, Johanne V Pastor, Animesh Nandi, Prem Gurnani, Owen P McGuinness, Hirotaka Chikuda, Masayuki Yamaguchi, Hiroshi Kawaguchi, Iichiro Shimomura, Yoshiharu Takayama, Joachim Herz, C Ronald Kahn, Kevin P Rosenblatt, and Makoto Kuro-o. Suppression of aging in mice by the hormone klotho. Science, 309(5742):1829–33, Sep 2005. doi: 10.1126/science.1112766.

[4] Dena B Dubal, Jennifer S Yokoyama, Lei Zhu, Lauren Broestl, Kurtresha Worden, Dan Wang, Virginia E Sturm, Daniel Kim, Eric Klein, Gui-Qiu Yu, Kaitlyn Ho, Kirsten E Eilertson, Lei Yu, Makoto Kuro-o, Philip L De Jager, Giovanni Coppola, Gary W Small, David A Bennett, Joel H Kramer, Carmela R Abraham, Bruce L Miller, and Lennart Mucke. Life extension factor klotho enhances cognition. Cell Rep, 7(4):1065–76, May 2014. doi: 10.1016/j.celrep.2014.03.076.

[5] Stacy A Castner, Shweta Gupta, Dan Wang, Arturo J Moreno, Cana Park, Chen Chen, Yan Poon, Aaron Groen, Kenneth Greenberg, Nathaniel David, Tom Boone, Mark G Baxter, Graham V Williams, and Dena B Dubal. Longevity factor klotho enhances cognition in aged nonhuman primates. Nat Aging, 3(8):931–937, Aug 2023. doi: 10.1038/s43587-023-00441-x.

[6] Yue Zhao, Chen-Ye Zeng, Xiao-Hong Li, Ting-Ting Yang, Xi Kuang, and Jun-Rong Du. Klotho overexpression improves amyloid-beta clearance and cognition in the APP/PS1 mouse model of Alzheimer’s disease. Aging Cell, 19(10):e13239, Sep 2020. doi: 10.1111/acel.13239.

[7] S T Sherry, M H Ward, M Kholodov, J Baker, L Phan, E M Smigielski, and K Sirotkin. dbsnp: the ncbi database of genetic variation. Nucleic Acids Res, 29(1):308–11, Jan 2001. doi: 10.1093/nar/29.1.308.

[8] Dan E Arking, Alice Krebsova, Milan Macek, Sr, Milan Macek, Jr, Albert Arking, I Saira Mian, Linda Fried, Ada Hamosh, Srabani Dey, Iain McIntosh, and Harry C Dietz. Association of human aging with a functional variant of klotho. Proc Natl Acad Sci U S A, 99(2):856–61, Jan 2002. doi: 10.1073/pnas.022484299.

[9] Danilo Di Bona, Giulia Accardi, Claudia Virruso, Giuseppina Candore, and Calogero Caruso. Association of klotho polymorphisms with healthy aging: a systematic review and meta-analysis. Rejuvenation Res, 17(2):212–6, Apr 2014. doi: 10.1089/rej.2013.1523.

[10] Dan E Arking, Gil Atzmon, Albert Arking, Nir Barzilai, and Harry C Dietz. Association between a functional variant of the KLOTHO gene and high-density lipoprotein cholesterol, blood pressure, stroke, and longevity. Circ Res, 96(4):412–8, Mar 2005. doi: 10.1161/01.RES.0000157171.04054.30.

[11] Laura Invidia, Stefano Salvioli, Serena Altilia, Michela Pierini, Maria P Panourgia, Daniela Monti, Francesco De Rango, Giuseppe Passarino, and Claudio Franceschi. The frequency of Klotho KL-VS polymorphism in a large italian population, from young subjects to centenarians, suggests the presence of specific time windows for its effect. Biogerontology, 11(1):67–73, Feb 2010. doi: 10.1007/s10522-009-9229-z.

[12] Zewu Zhu, Weiping Xia, Yu Cui, Feng Zeng, Yang Li, Zhongqing Yang, and Chen Hequn. Klotho gene polymorphisms are associated with healthy aging and longevity: Evidence from a meta-analysis. Mech Ageing Dev, 178:33–40, Mar 2019. doi: 10.1016/j.mad.2018.12.003.

[13] M V Garcia, J A Cabezas, and M N Perez-Gonzalez. Alterations in the activities of subcellular fractions marker enzymes in rat liver and brain by hydrocortisone and corticosterone treatment. Int J Biochem, 17(2):203–8, 1985. doi: 10.1016/0020-711x(85)90115-6.

[14] Jennifer S Yokoyama, Virginia E Sturm, Luke W Bonham, Eric Klein, Konstantinos Arfanakis, Lei Yu, Giovanni Coppola, Joel H Kramer, David A Bennett, Bruce L Miller, and Dena B Dubal. Variation in longevity gene KLOTHO is associated with greater cortical volumes. Ann Clin Transl Neurol, 2(3): 215–30, Mar 2015. doi: 10.1002/acn3.161.

[15] Michael E Belloy, Valerio Napolioni, Summer S Han, Yann Le Guen, Michael D Greicius, and Alzheimer’s Disease Neuroimaging Initiative. Association of Klotho-VS heterozygosity with risk of Alzheimer disease in individuals who carry APOE4. JAMA Neurol, 77(7):849–862, Jul 2020. doi: 10.1001/jamaneurol.2020.0414.

[16] Claire M Erickson, Stephanie A Schultz, Jennifer M Oh, Burcu F Darst, Yue Ma, Derek Norton, Tobey Betthauser, Catherine L Gallagher, Cynthia M Carlsson, Barbara B Bendlin, Sanjay Asthana, Bruce P Hermann, Mark A Sager, Kaj Blennow, Henrik Zetterberg, Corinne D Engelman, Bradley T Christian, Sterling C Johnson, Dena B Dubal, and Ozioma C Okonkwo. KLOTHO heterozygosity attenuates APOE4-related amyloid burden in preclinical AD. Neurology, 92(16):e1878–e1889, Apr 2019. doi: 10.1212/WNL.0000000000007323.

[17] Nijee S Luthra, Luke W Bonham, Arturo J Moreno, Cana Park, Claire J C Huguenard, Samira Abdulai-Saiku, Shweta Gupta, Jonathan Lin, Lauren Broestl, Alexandre Bétourné, Rohan Sehgal, Dan Wang, Sylvain E Lesné, Jill L Ostrem, Jennifer S Yokoyama, and Dena B Dubal. Longevity factor klotho and resistance to cognitive deficits in individuals with parkinson’s disease and in an -synuclein mouse model. J Neurosci, page e1904252026, Mar 2026. doi: 10.1523/JNEUROSCI.1904-25.2026.

[18] Alessandra Errigo, Maria Pina Dore, Giammarco Mocci, and Giovanni Mario Pes. Lack of association between common polymorphisms associated with successful aging and longevity in the population of Sardinian Blue Zone. Sci Rep, 14(1):30773, Dec 2024. doi: 10.1038/s41598-024-80497-w.

[19] Clarisse F de Vries, Roger T Staff, Kimberly G Noble, Ryan L Muetzel, Meike W Vernooij, Tonya White, Gordon D Waiter, Alison D Murray, and Pediatric Imaging, Neurocognition and Genetics Study. Klotho gene polymorphism, brain structure and cognition in early-life development. Brain Imaging Behav, 14 (1):213–225, Feb 2020. doi: 10.1007/s11682-018-9990-1.

[20] Kengo Shibata, Cheng Chen, Xin You Tai, Sanjay G Manohar, and Masud Husain. Impact of apoe, klotho, and sex on cognitive decline with aging. Proc Natl Acad Sci U S A, 122(6):e2416042122, Feb 2025. doi: 10.1073/pnas.2416042122.

[21] Dan E Arking, Diane M Becker, Lisa R Yanek, Daniele Fallin, Daniel P Judge, Taryn F Moy, Lewis C Becker, and Harry C Dietz. KLOTHO allele status and the risk of early-onset occult coronary artery disease. Am J Hum Genet, 72(5):1154–61, May 2003. doi: 10.1086/375035.

[22] Osvaldo P Almeida, Bharti Morar, Graeme J Hankey, Bu B Yeap, Jonathan Golledge, Assen Jablensky, and Leon Flicker. Longevity Klotho gene polymorphism and the risk of dementia in older men. Maturitas, 101:1–5, Jul 2017. doi: 10.1016/j.maturitas.2017.04.005.

[23] Si Liu, Mingyang Wu, Yan Wang, Lu Xiang, Gang Luo, Qian Lin, and Lin Xiao. The association between dietary fiber intake and serum klotho levels in americans: A cross-sectional study from the national health and nutrition examination survey. Nutrients, 15(14):3147, Jul 2023. doi: 10.3390/nu15143147.

[24] Gisele Pereira Dias, Tytus Murphy, Doris Stangl, Selda Ahmet, Benjamin Morisse, Alina Nix, Lindsey J Aimone, James B Aimone, Makoto Kuro-O, Fred H Gage, and Sandrine Thuret. Intermittent fasting enhances long-term memory consolidation, adult hippocampal neurogenesis, and expression of longevity gene klotho. Mol Psychiatry, 26(11):6365–6379, Nov 2021. doi: 10.1038/s41380-021-01102-4.

[25] Tomoko Matsubara, Asako Miyaki, Nobuhiko Akazawa, Youngju Choi, Song-Gyu Ra, Koichiro Tanahashi, Hiroshi Kumagai, Satoshi Oikawa, and Seiji Maeda. Aerobic exercise training increases plasma klotho levels and reduces arterial stiffness in postmenopausal women. Am J Physiol Heart Circ Physiol, 306(3):H348–55, Feb 2014. doi: 10.1152/ajpheart.00429.2013.

[26] A A Prather, E S Epel, J Arenander, L Broestl, B I Garay, D Wang, and D B Dubal. Longevity factor klotho and chronic psychological stress. Transl Psychiatry, 5(6):e585, Jun 2015. doi: 10.1038/tp.2015.81.

[27] Thomas O Carpenter, Karl L Insogna, Jane H Zhang, Bruce Ellis, Sherril Nieman, Christine Simpson, Elizabeth Olear, and Caren M Gundberg. Circulating levels of soluble klotho and fgf23 in x-linked hypophosphatemia: circadian variance, effects of treatment, and relationship to parathyroid status. J Clin Endocrinol Metab, 95(11):E352–7, Nov 2010. doi: 10.1210/jc.2010-0589.

[28] Michael I Love, Wolfgang Huber, and Simon Anders. Moderated estimation of fold change and dispersion for rna-seq data with deseq2. Genome Biol, 15(12):550, 2014. doi: 10.1186/s13059-014-0550-8.

[29] Malgorzata Nowicka and Mark D. Robinson. Drimseq: a dirichlet-multinomial framework for multivariate count outcomes in genomics [version 2; referees: 2 approved]. F1000Research, 5(1356), 2016. doi: 10.12688/f1000research.8900.2. URL https://f1000research.com/articles/5-1356/v2.

[30] Gregory A Cary, Jesse C Wiley, Jake Gockley, Stephen Keegan, Sai Sruthi Amirtha Ganesh, Laura Heath, Robert R Butler, 3rd, Lara M Mangravite, Benjamin A Logsdon, Frank M Longo, Allan Levey, Anna K Greenwood, and Gregory W Carter. Genetic and multi-omic risk assessment of Alzheimer’s disease implicates core associated biological domains. Alzheimers Dement (N Y), 10(2):e12461, 2024. doi: 10.1002/trc2.12461.

[31] Stephen Keegan, Gregory A. Cary, Jesse C Wiley, Karina Leal, Robert R. Butler III, Anna Greenwood, Gregory W Carter, and The Emory-Sage-SGC TREAT-AD Center. Organizing Alzheimer’s disease endophenotypes into subdivisions of biological domains improves therapeutic target hypothesis selection. Alzheimer’s & Dementia, 20(S1):e092649, 2024. doi: 10.1002/alz.092649. URL https://alz-journals.onlinelibrary.wiley.com/doi/abs/10.1002/alz.092649.

[32] M Kanehisa and S Goto. KEGG: Kyoto encyclopedia of genes and genomes. Nucleic Acids Res, 28(1): 27–30, Jan 2000. doi: 10.1093/nar/28.1.27.

[33] Minoru Kanehisa, Miho Furumichi, Yoko Sato, Masayuki Kawashima, and Mari Ishiguro-Watanabe. KEGG for taxonomy-based analysis of pathways and genomes. Nucleic Acids Res, 51(D1):D587–D592, Jan 2023. doi: 10.1093/nar/gkac963.

[34] Guangchuang Yu, Li-Gen Wang, Yanyan Han, and Qing-Yu He. clusterProfiler: an R package for comparing biological themes among gene clusters. OMICS, 16(5):284–7, May 2012. doi: 10.1089/omi.2011.0118.

[35] Steffen Durinck, Paul T Spellman, Ewan Birney, and Wolfgang Huber. Mapping identifiers for the integration of genomic datasets with the r/bioconductor package biomart. Nat Protoc, 4(8):1184–91, 2009. doi: 10.1038/nprot.2009.97.

[36] Steffen Durinck, Yves Moreau, Arek Kasprzyk, Sean Davis, Bart De Moor, Alvis Brazma, and Wolfgang Huber. Biomart and bioconductor: a powerful link between biological databases and microarray data analysis. Bioinformatics, 21(16):3439–40, Aug 2005. doi: 10.1093/bioinformatics/bti525.

[37] Mohammed R Shaker, Julio Aguado, Harman Kaur Chaggar, and Ernst J Wolvetang. Klotho inhibits neuronal senescence in human brain organoids. NPJ Aging Mech Dis, 7(1):18, Aug 2021. doi: 10.1038/s41514-021-00070-x.

[38] Tanya Barrett, Stephen E Wilhite, Pierre Ledoux, Carlos Evangelista, Irene F Kim, Maxim Toma-shevsky, Kimberly A Marshall, Katherine H Phillippy, Patti M Sherman, Michelle Holko, Andrey Yefanov, Hyeseung Lee, Naigong Zhang, Cynthia L Robertson, Nadezhda Serova, Sean Davis, and Alexandra Soboleva. Ncbi geo: archive for functional genomics data sets–update. Nucleic Acids Res, 41 (Database issue):D991–5, Jan 2013. doi: 10.1093/nar/gks1193.

[39] Ron Edgar, Michael Domrachev, and Alex E Lash. Gene expression omnibus: Ncbi gene expression and hybridization array data repository. Nucleic Acids Res, 30(1):207–10, Jan 2002. doi: 10.1093/nar/30.1.207.

[40] Jennifer S Yokoyama, Gabe Marx, Jesse A Brown, Luke W Bonham, Dan Wang, Giovanni Coppola, William W Seeley, Howard J Rosen, Bruce L Miller, Joel H Kramer, and Dena B Dubal. Systemic klotho is associated with KLOTHO variation and predicts intrinsic cortical connectivity in healthy human aging. Brain Imaging Behav, 11(2):391–400, Apr 2017. doi: 10.1007/s11682-016-9598-2.

[41] Julian M Gaitán, Sanjay Asthana, Cynthia M Carlsson, Corinne D Engelman, Sterling C Johnson, Mark A Sager, Dan Wang, Dena B Dubal, and Ozioma C Okonkwo. Circulating Klotho is higher in cerebrospinal fluid than serum and elevated among KLOTHO heterozygotes in a cohort with risk for Alzheimer’s disease. J Alzheimers Dis, 90(4):1557–1569, 2022. doi: 10.3233/JAD-220571.

[42] Ian J Deary, Sarah E Harris, Helen C Fox, Caroline Hayward, Alan F Wright, John M Starr, and Lawrence J Whalley. Klotho genotype and cognitive ability in childhood and old age in the same individuals. Neurosci Lett, 378(1):22–7, Apr 2005. doi: 10.1016/j.neulet.2004.12.005.

[43] C. Behl. Alzheimer’s Disease Research: What Has Guided Research So Far and Why It Is High Time for a Paradigm Shift. Biomedical and Life Sciences. Springer International Publishing, 2023. ISBN 9783031315701. URL https://books.google.com/books?id=EnPLEAAAQBAJ.

[44] Lisa Pham, Lisa Christadore, Scott Schaus, and Eric D Kolaczyk. Network-based prediction for sources of transcriptional dysregulation using latent pathway identification analysis. Proc Natl Acad Sci U S A, 108(32):13347–52, Aug 2011. doi: 10.1073/pnas.1100891108.

[45] Zheng Jia, Qian Liu, Ying Xie, Jie Wei, Xiaolin Sun, Fandi Meng, Bin Zhao, Zhenkun Yu, Li Zhao, and Zhengjiang Xing. Klotho/FGF23 axis regulates cardiomyocyte apoptosis and cytokine release through ERK/MAPK pathway. Cardiovasc Toxicol, 23(9-10):317–328, Oct 2023. doi: 10.1007/s12012-023-09805-6.

[46] Taslim Uddin. Oxidative genomic or genotoxic stress in neurodegeneration: Mechanisms and therapeutic avenues. AIMS Neurosci, 12(3):369–390, 2025. doi: 10.3934/Neuroscience.2025020.

[47] Ju Gao, Lauren Vicuna, and Xinglong Wang. Organelle abnormalities in alzheimer’s disease. Organelle, 3, 2025. doi: 10.61747/0ifp.202503005.

[48] Michael T Lin and M Flint Beal. Mitochondrial dysfunction and oxidative stress in neurodegenerative diseases. Nature, 443(7113):787–95, Oct 2006. doi: 10.1038/nature05292.

[49] Wenzhang Wang, Fanpeng Zhao, Xiaopin Ma, George Perry, and Xiongwei Zhu. Mitochondria dysfunction in the pathogenesis of alzheimer’s disease: recent advances. Mol Neurodegener, 15(1):30, May 2020. doi: 10.1186/s13024-020-00376-6.

[50] Cindy V Ly and Patrik Verstreken. Mitochondria at the synapse. Neuroscientist, 12(4):291–9, Aug 2006. doi: 10.1177/1073858406287661.

[51] Yasunori Hayashi. Molecular mechanism of hippocampal long-term potentiation - towards multiscale understanding of learning and memory. Neurosci Res, 175:3–15, Feb 2022. doi: 10.1016/j.neures.2021.08.001.

[52] Han-Jun Wu, Wen-Ning Wu, Hua Fan, Liu-Er Liu, Jin-Qiong Zhan, Yi-Heng Li, Chun-Nuan Chen, Shu-Zhen Jiang, Jian-Wen Xiong, Zhi-Min Yu, Bo Wei, Wei Wang, and Yuan-Jian Yang. Life extension factor klotho regulates behavioral responses to stress via modulation of glun2b function in the nucleus accumbens. Neuropsychopharmacology, 47(9):1710–1720, Aug 2022. doi: 10.1038/s41386-022-01323-3.

[53] Ella Zeldich, Ci-Di Chen, Teresa A Colvin, Erin A Bove-Fenderson, Jennifer Liang, Tracey B Tucker Zhou, David A Harris, and Carmela R Abraham. The neuroprotective effect of klotho is mediated via regulation of members of the redox system. J Biol Chem, 289(35):24700–15, Aug 2014. doi: 10.1074/jbc.M114.567321.

[54] Dena B Dubal, Lei Zhu, Pascal E Sanchez, Kurtresha Worden, Lauren Broestl, Erik Johnson, Kaitlyn Ho, Gui-Qiu Yu, Daniel Kim, Alexander Betourne, Makoto Kuro-O, Eliezer Masliah, Carmela R Abraham, and Lennart Mucke. Life extension factor klotho prevents mortality and enhances cognition in happ transgenic mice. J Neurosci, 35(6):2358–71, Feb 2015. doi: 10.1523/JNEUROSCI.5791-12.2015.

[55] Cana Park, Oliver Hahn, Shweta Gupta, Arturo J Moreno, Francesca Marino, Blen Kedir, Dan Wang, Saul A Villeda, Tony Wyss-Coray, and Dena B Dubal. Platelet factors are induced by longevity factor klotho and enhance cognition in young and aging mice. Nat Aging, 3(9):1067–1078, Sep 2023. doi: 10.1038/s43587-023-00468-0.

[56] Foluwasomi A Oyefeso, Alysson R Muotri, Christopher G Wilson, and Michael J Pecaut. Brain organoids: A promising model to assess oxidative stress-induced central nervous system damage. Dev Neurobiol, 81(5):653–670, Jul 2021. doi: 10.1002/dneu.22828.

[57] Michael E Belloy, Sarah J Eger, Yann Le Guen, Valerio Napolioni, Kacie D Deters, Hyun-Sik Yang, Marzia A Scelsi, Tenielle Porter, Sarah-Naomi James, Andrew Wong, Jonathan M Schott, Reisa A Sperling, Simon M Laws, Elisabeth C Mormino, Zihuai He, Summer S Han, Andre Altmann, Michael D Greicius, A4 Study Team, Insight 46 Study Team, Australian Imaging Biomarkers and Lifestyle (AIBL) Study, and Alzheimer’s Disease Neuroimaging Initiative. KL-VS heterozygosity reduces brain amyloid in asymptomatic at-risk APOE4 carriers. Neurobiol Aging, 101:123–129, May 2021. doi: 10.1016/j.neurobiolaging.2021.01.008.

[58] Xinglong Wang, Bo Su, Hyoung-gon Lee, Xinyi Li, George Perry, Mark A Smith, and Xiongwei Zhu. Impaired balance of mitochondrial fission and fusion in alzheimer’s disease. J Neurosci, 29(28):9090–103, Jul 2009. doi: 10.1523/JNEUROSCI.1357-09.2009.

[59] Venkatesh N Murthy and Pietro De Camilli. Cell biology of the presynaptic terminal. Annu Rev Neurosci, 26:701–28, 2003. doi: 10.1146/annurev.neuro.26.041002.131445.

[60] R Rizzuto. Intracellular ca(2+) pools in neuronal signalling. Curr Opin Neurobiol, 11(3):306–11, Jun 2001. doi: 10.1016/s0959-4388(00)00212-9.

[61] Rosario Rizzuto, Michael R Duchen, and Tullio Pozzan. Flirting in little space: the er/mitochondria ca2+ liaison. Sci STKE, 2004(215):re1, Jan 2004. doi: 10.1126/stke.2152004re1.

[62] Qunxing Ding, William R Markesbery, Qinghua Chen, Feng Li, and Jeffrey N Keller. Ribosome dysfunction is an early event in alzheimer’s disease. J Neurosci, 25(40):9171–5, Oct 2005. doi: 10.1523/JNEUROSCI.3040-05.2005.

[63] Sebastian Iben. To aggregate or not to aggregate - is it a matter of the ribosome? Bioessays, 45(7): e2200230, Jul 2023. doi: 10.1002/bies.202200230.

[64] Jack Llewellyn, Venkatesh Mallikarjun, Ellen Appleton, Maria Osipova, Hamish T J Gilbert, Stephen M Richardson, Simon J Hubbard, and Joe Swift. Loss of regulation of protein synthesis and turnover underpins an attenuated stress response in senescent human mesenchymal stem cells. Proc Natl Acad Sci U S A, 120(14):e2210745120, Apr 2023. doi: 10.1073/pnas.2210745120.

[65] Margarita Brilkova, Martina Nigri, Harshitha Santhosh Kumar, James Moore, Matilde Mantovani, Claudia Keller, Amandine Grimm, Anne Eckert, Dimitri Shcherbakov, Rashid Akbergenov, Petra Seebeck, Stefanie D Krämer, David P Wolfer, Thomas C Gent, and Erik C Böttger. Error-prone protein synthesis recapitulates early symptoms of alzheimer disease in aging mice. Cell Rep, 40(13):111433, Sep 2022. doi: 10.1016/j.celrep.2022.111433.

[66] Yoo Jin Shin, Sun Woo Lim, Sheng Cui, Eun Jeong Ko, Byung Ha Chung, Hong Lim Kim, Tae Ryong Riew, Mun Yong Lee, and Chul Woo Yang. Tacrolimus decreases cognitive function by impairing hippocampal synaptic balance: a possible role of klotho. Mol Neurobiol, 58(11):5954–5970, Nov 2021. doi: 10.1007/s12035-021-02499-3.

[67] Eugene V Mosharov, Ayelet M Rosenberg, Anna S Monzel, Corey A Osto, Linsey Stiles, Gorazd B Rosoklija, Andrew J Dwork, Snehal Bindra, Alex Junker, Ya Zhang, Masashi Fujita, Madeline B Mariani, Mihran Bakalian, David Sulzer, Philip L De Jager, Vilas Menon, Orian S Shirihai, J John Mann, Mark D Underwood, Maura Boldrini, Michel Thiebaut de Schotten, and Martin Picard. A human brain map of mitochondrial respiratory capacity and diversity. Nature, 641(8063):749–758, May 2025. doi: 10.1038/s41586-025-08740-6.

[68] A S Stoykova, M D Dabeva, R N Dimova, and A A Hadjiolov. Ribosome biogenesis and nucleolar ultrastructure in neuronal and oligodendroglial rat brain cells. J Neurochem, 45(6):1667–76, Dec 1985. doi: 10.1111/j.1471-4159.1985.tb10521.x.

[69] Yingxue Ren, Xue Wang, Shuwen Zhang, Hongru Hu, Zachary Quicksall, Sangderk Lee, Josh M Morganti, Lance A Johnson, Yan W Asmann, and Na Zhao. Deconvolution reveals cell-type-specific transcriptomic changes in the aging mouse brain. Sci Rep, 13(1):16855, Oct 2023. doi: 10.1038/s41598-023-44183-7.

[70] Methodios Ximerakis, Scott L Lipnick, Brendan T Innes, Sean K Simmons, Xian Adiconis, Danielle Dionne, Brittany A Mayweather, Lan Nguyen, Zachary Niziolek, Ceren Ozek, Vincent L Butty, Ruth Isserlin, Sean M Buchanan, Stuart S Levine, Aviv Regev, Gary D Bader, Joshua Z Levin, and Lee L Rubin. Single-cell transcriptomic profiling of the aging mouse brain. Nat Neurosci, 22(10):1696–1708, Oct 2019. doi: 10.1038/s41593-019-0491-3.

[71] Kelly Jin, Zizhen Yao, Cindy T J van Velthoven, Eitan S Kaplan, Katie Glattfelder, Samuel T Barlow, Gabriella Boyer, Daniel Carey, Tamara Casper, Anish Bhaswanth Chakka, Rushil Chakrabarty, Michael Clark, Max Departee, Marie Desierto, Amanda Gary, Jessica Gloe, Jeff Goldy, Nathan Guilford, Junitta Guzman, Daniel Hirschstein, Changkyu Lee, Elizabeth Liang, Trangthanh Pham, Melissa Reding, Kara Ronellenfitch, Augustin Ruiz, Josh Sevigny, Nadiya Shapovalova, Lyudmila Shulga, Josef Sulc, Amy Torkelson, Herman Tung, Boaz Levi, Susan M Sunkin, Nick Dee, Luke Esposito, Kimberly A Smith, Bosiljka Tasic, and Hongkui Zeng. Brain-wide cell-type-specific transcriptomic signatures of healthy ageing in mice. Nature, 638(8049):182–196, Feb 2025. doi: 10.1038/s41586-024-08350-8.

[72] Céline Bellenguez, Fahri Küçükali, Iris E Jansen, Luca Kleineidam, Sonia Moreno-Grau, Najaf Amin, Adam C Naj, Rafael Campos-Martin, Benjamin Grenier-Boley, Victor Andrade, Peter A Holmans, Anne Boland, Vincent Damotte, Sven J van der Lee, Marcos R Costa, Teemu Kuulasmaa, Qiong Yang, Itziar de Rojas, Joshua C Bis, Amber Yaqub, Ivana Prokic, Julien Chapuis, Shahzad Ahmad, Vilmantas Giedraitis, Dag Aarsland, Pablo Garcia-Gonzalez, Carla Abdelnour, Emilio Alarcón-Martín, Daniel Alcolea, Montserrat Alegret, Ignacio Alvarez, Victoria Álvarez, Nicola J Armstrong, Anthoula Tsolaki, Carmen Antúnez, Ildebrando Appollonio, Marina Arcaro, Silvana Archetti, Alfonso Arias Pastor, Beatrice Arosio, Lavinia Athanasiu, Henri Bailly, Nerisa Banaj, Miquel Baquero, Sandra Barral, Alexa Beiser, Ana Belén Pastor, Jennifer E Below, Penelope Benchek, Luisa Benussi, Claudine Berr, Céline Besse, Valentina Bessi, Giuliano Binetti, Alessandra Bizarro, Rafael Blesa, Mercè Boada, Eric Boerwinkle, Barbara Borroni, Silvia Boschi, Paola Bossù, Geir Bråthen, Jan Bressler, Catherine Bres-ner, Henry Brodaty, Keeley J Brookes, Luis Ignacio Brusco, Dolores Buiza-Rueda, Katharina Bûrger, Vanessa Burholt, William S Bush, Miguel Calero, Laura B Cantwell, Geneviève Chene, Jaeyoon Chung, Michael L Cuccaro, Ángel Carracedo, Roberta Cecchetti, Laura Cervera-Carles, Camille Charbonnier, Hung-Hsin Chen, Caterina Chillotti, Simona Ciccone, Jurgen A H R Claassen, Christopher Clark, Elisa Conti, Anaïs Corma-Gómez, Emanuele Costantini, Carlo Custodero, Delphine Daian, Maria Carolina Dalmasso, Antonio Daniele, Efthimios Dardiotis, Jean-François Dartigues, Peter Paul de Deyn, Ka-tia de Paiva Lopes, Lot D de Witte, Stéphanie Debette, Jürgen Deckert, Teodoro Del Ser, Nicola Denning, Anita DeStefano, Martin Dichgans, Janine Diehl-Schmid, Mónica Diez-Fairen, Paolo Dionigi Rossi, Srdjan Djurovic, Emmanuelle Duron, Emrah Düzel, Carole Dufouil, Gudny Eiriksdottir, Sebas-tiaan Engelborghs, Valentina Escott-Price, Ana Espinosa, Michael Ewers, Kelley M Faber, Tagliavini Fabrizio, Sune Fallgaard Nielsen, David W Fardo, Lucia Farotti, Chiara Fenoglio, Marta Fernández-Fuertes, Raffaele Ferrari, Catarina B Ferreira, Evelyn Ferri, Bertrand Fin, Peter Fischer, Tormod Fladby, Klaus Fließbach, Bernard Fongang, Myriam Fornage, Juan Fortea, Tatiana M Foroud, Sil-via Fostinelli, Nick C Fox, Emlio Franco-Macías, María J Bullido, Ana Frank-García, Lutz Froelich, Brian Fulton-Howard, Daniela Galimberti, Jose Maria García-Alberca, Pablo García-González, Sebas-tian Garcia-Madrona, Guillermo Garcia-Ribas, Roberta Ghidoni, Ina Giegling, Giaccone Giorgio, Ali-son M Goate, Oliver Goldhardt, Duber Gomez-Fonseca, Antonio González-Pérez, Caroline Graff, Giulia Grande, Emma Green, Timo Grimmer, Edna Grünblatt, Michelle Grunin, Vilmundur Gudnason, Tamar Guetta-Baranes, Annakaisa Haapasalo, Georgios Hadjigeorgiou, Jonathan L Haines, Kara L Hamilton-Nelson, Harald Hampel, Olivier Hanon, John Hardy, Annette M Hartmann, Lucrezia Hausner, Janet Harwood, Stefanie Heilmann-Heimbach, Seppo Helisalmi, Michael T Heneka, Isabel Hernández, Mar-tin J Herrmann, Per Hoffmann, Clive Holmes, Henne Holstege, Raquel Huerto Vilas, Marc Hulsman, Jack Humphrey, Geert Jan Biessels, Xueqiu Jian, Charlotte Johansson, Gyungah R Jun, Yuriko Kas-tumata, John Kauwe, Patrick G Kehoe, Lena Kilander, Anne Kinhult Ståhlbom, Miia Kivipelto, Anne Koivisto, Johannes Kornhuber, Mary H Kosmidis, Walter A Kukull, Pavel P Kuksa, Brian W Kunkle, Amanda B Kuzma, Carmen Lage, Erika J Laukka, Lenore Launer, Alessandra Lauria, Chien-Yueh Lee, Jenni Lehtisalo, Ondrej Lerch, Alberto Lleó, William Longstreth, Jr, Oscar Lopez, Adolfo Lopez de Munain, Seth Love, Malin Löwemark, Lauren Luckcuck, Kathryn L Lunetta, Yiyi Ma, Juan Macías, Catherine A MacLeod, Wolfgang Maier, Francesca Mangialasche, Marco Spallazzi, Marta Marquié, Rachel Marshall, Eden R Martin, Angel Martín Montes, Carmen Martínez Rodríguez, Carlo Masullo, Richard Mayeux, Simon Mead, Patrizia Mecocci, Miguel Medina, Alun Meggy, Shima Mehrabian, Silvia Mendoza, Manuel Menéndez-González, Pablo Mir, Susanne Moebus, Merel Mol, Laura Molina-Porcel, Laura Montrreal, Laura Morelli, Fermin Moreno, Kevin Morgan, Thomas Mosley, Markus M Nöthen, Carolina Muchnik, Shubhabrata Mukherjee, Benedetta Nacmias, Tiia Ngandu, Gael Nicolas, Børge G Nordestgaard, Robert Olaso, Adelina Orellana, Michela Orsini, Gemma Ortega, Alessandro Padovani, Caffarra Paolo, Goran Papenberg, Lucilla Parnetti, Florence Pasquier, Pau Pastor, Gina Peloso, Alba Pérez-Cordón, Jordi Pérez-Tur, Pierre Pericard, Oliver Peters, Yolande A L Pijnenburg, Juan A Pineda, Gerard Piñol-Ripoll, Claudia Pisanu, Thomas Polak, Julius Popp, Danielle Posthuma, Josef Priller, Raquel Puerta, Olivier Quenez, Inés Quintela, Jesper Qvist Thomassen, Alberto Rábano, Innocenzo Rainero, Farid Rajabli, Inez Ramakers, Luis M Real, Marcel J T Reinders, Christiane Reitz, Dolly Reyes-Dumeyer, Perry Ridge, Steffi Riedel-Heller, Peter Riederer, Natalia Roberto, Eloy Rodriguez-Rodriguez, Arvid Rongve, Irene Rosas Allende, Maitée Rosende-Roca, Jose Luis Royo, Elisa Rubino, Dan Rujescu, María Eugenia Sáez, Paraskevi Sakka, Ingvild Saltvedt, Ángela Sanabria, María Bernal Sánchez-Arjona, Florentino Sanchez-Garcia, Pascual Sánchez Juan, Raquel Sánchez-Valle, Sigrid B Sando, Chloé Sarnowski, Claudia L Satizabal, Michela Scamosci, Nikolaos Scarmeas, Elio Scarpini, Philip Scheltens, Norbert Scherbaum, Martin Scherer, Matthias Schmid, Anja Schneider, Jonathan M Schott, Geir Selbæk, Davide Seripa, Manuel Serrano, Jin Sha, Alexey A Shadrin, Olivia Skrobot, Susan Slifer, Gijsje J L Snijders, Hilkka Soininen, Vincenzo Solfrizzi, Alina Solomon, Yeunjoo Song, Sandro Sorbi, Oscar Sotolongo-Grau, Gianfranco Spalletta, Annika Spottke, Alessio Squassina, Eystein Stordal, Juan Pablo Tartan, Lluís Tárraga, Niccolo Tesí, Anbupalam Thalamuthu, Tegos Thomas, Giuseppe Tosto, Latchezar Traykov, Lucio Tremolizzo, Anne Tybjærg-Hansen, Andre Uitterlinden, Abbe Ullgren, Ingun Ulstein, Sergi Valero, Otto Valladares, Christine Van Broeckhoven, Jeffery Vance, Badri N Var-darajan, Aad van der Lugt, Jasper Van Dongen, Jeroen van Rooij, John van Swieten, Rik Vandenberghe, Frans Verhey, Jean-Sébastien Vidal, Jonathan Vogelgsang, Martin Vyhnalek, Michael Wagner, David Wallon, Li-San Wang, Ruiqi Wang, Leonie Weinhold, Jens Wiltfang, Gill Windle, Bob Woods, Mary Yannakoulia, Habil Zare, Yi Zhao, Xiaoling Zhang, Congcong Zhu, Miren Zulaica, EADB, GR@ACE, DEGESCO, EADI, GERAD, Demgene, FinnGen, ADGC, CHARGE, Lindsay A Farrer, Bruce M Psaty, Mohsen Ghanbari, Towfique Raj, Perminder Sachdev, Karen Mather, Frank Jessen, M Arfan Ikram, Alexandre de Mendonça, Jakub Hort, Magda Tsolaki, Margaret A Pericak-Vance, Philippe Amouyel, Julie Williams, Ruth Frikke-Schmidt, Jordi Clarimon, Jean-François Deleuze, Giacomina Rossi, Sudha Seshadri, Ole A Andreassen, Martin Ingelsson, Mikko Hiltunen, Kristel Sleegers, Gerard D Schellen-berg, Cornelia M van Duijn, Rebecca Sims, Wiesje M van der Flier, Agustín Ruiz, Alfredo Ramirez, and Jean-Charles Lambert. New insights into the genetic etiology of alzheimer’s disease and related dementias. Nat Genet, 54(4):412–436, Apr 2022. doi: 10.1038/s41588-022-01024-z.

[73] Oleg Butovsky, Neta Rosenzweig, Kilian L Kleemann, Mehdi Jorfi, Vijay K Kuchroo, Rudolph E Tanzi, and Howard L Weiner. Immune dysfunction in alzheimer disease. Nat Rev Neurosci, Nov 2025. doi: 10.1038/s41583-025-00997-0.

[74] Zaw Myo Hein, Barani Karikalan, Prarthana Kalerammana Gopalakrishna, Krina Dhevi, Aisyah Alkatiri, Farida Hussan, Mohamad Aris Mohd Moklas, Saravanan Jagadeesan, Muhammad Danial Che Ramli, Che Mohd Nasril Che Mohd Nassir, and Thirupathirao Vishnumukkala. Toward a unified framework in molecular neurobiology of alzheimer’s disease: Revisiting the pathophysiological hypotheses. Mol Neurobiol, 63(1):282, Dec 2025. doi: 10.1007/s12035-025-05602-0.

[75] Fangda Leng and Paul Edison. Neuroinflammation and microglial activation in alzheimer disease: where do we go from here? Nat Rev Neurol, 17(3):157–172, Mar 2021. doi: 10.1038/s41582-020-00435-y.

[76] Hadas Keren-Shaul, Amit Spinrad, Assaf Weiner, Orit Matcovitch-Natan, Raz Dvir-Szternfeld, Tyler K Ulland, Eyal David, Kuti Baruch, David Lara-Astaiso, Beata Toth, Shalev Itzkovitz, Marco Colonna, Michal Schwartz, and Ido Amit. A unique microglia type associated with restricting development of alzheimer’s disease. Cell, 169(7):1276–1290.e17, Jun 2017. doi: 10.1016/j.cell.2017.05.018.

[77] Meng-Shan Tan, Jin-Tai Yu, Teng Jiang, Xi-Chen Zhu, and Lan Tan. The nlrp3 inflammasome in alzheimer’s disease. Mol Neurobiol, 48(3):875–82, Dec 2013. doi: 10.1007/s12035-013-8475-x.

[78] Xi Richard Chen, Yongzhao Shao, Martin J Sadowski, and On Behalf Of The Alzheimer’s Disease Neuroimaging Initiative. Interaction between KLOTHO-VS heterozygosity and APOE E4 allele predicts rate of cognitive decline in late-onset Alzheimer’s disease. Genes (Basel), 14(4):917, Apr 2023. doi: 10.3390/genes14040917.

